# The Deubiquitylase Ubp15 Couples Transcription to mRNA Export

**DOI:** 10.1101/2020.08.20.258822

**Authors:** Fanny Eyboulet, Célia Jeronimo, Jacques Côté, François Robert

## Abstract

The nuclear export of messenger RNAs (mRNAs) is intimately coupled to their synthesis. pre-mRNAs assemble into dynamic ribonucleoparticles as they are being transcribed, processed and exported. The role of ubiquitylation in this process is increasingly recognized as the ubiquitylation of many key players have been shown to affect mRNA nuclear export. While a few E3 ligases have been shown to regulate nuclear export, evidence for deubiquitylases is currently lacking. Here, we identified the deubiquitylase Ubp15 as a regulator of nuclear export in *Saccharomyces cerevisiae*. Ubp15 interacts both with RNA polymerase II and with the nuclear pore complex, and its deletion reverts the nuclear export defect of mutants of the E3 ligase Rsp5. The deletion of *UBP15* leads to hyper-ubiquitylation of the main nuclear export receptor Mex67 and affects its association with THO, a complex coupling transcription to mRNA processing and involved in the recruitment of mRNA export factors to nascent transcripts. Collectively, our data support a role for Ubp15 in coupling transcription to mRNA export.

## INTRODUCTION

In eukaryotes, RNA polymerase II (RNAPII) is the enzyme responsible for the transcription of all protein-coding genes, as well as several noncoding RNAs. Rpb1, the largest subunit of RNAPII, contains a C-terminal domain (CTD) composed of 26 to 52 repetitions (in yeast and mammals respectively) of the heptapeptide Y_1_S_2_P_3_T_4_S_5_P_6_S_7_ (Chapman *et al*., 2008). The CTD is highly conserved and essential for viability in all organisms. During transcription, the CTD is extensively and dynamically phosphorylated to coordinate the binding of proteins involved in the different steps of transcription and to couple transcription to mRNA processing (Corden, 2013, Eick *et al*., 2013, Harlen *et al*., 2017, Jeronimo *et al*., 2013, Jeronimo *et al*., 2016).

During their synthesis, processing and export, pre-mRNAs and mature mRNAs are packaged with RNA-binding proteins to form dynamic ribonucleoprotein particles (mRNPs) (Mitchell *et al*., 2014, Singh *et al*., 2015, Tutucci *et al*., 2011). The mRNA export machinery is highly conserved from yeast to humans and mainly depends on the export factor Mex67-Mtr2 (TAP-NXF1 in mammals) (Nino *et al*., 2013, Scott *et al*., 2019, Wende *et al*., 2019). Mex67, in complex with Mtr2, mediates transport through the nuclear pore complex (NPC) via interactions with several FG-enriched nucleoporins (Nups) (Santos-Rosa *et al*., 1998, Strasser *et al*., 2000a, Terry *et al*., 2007). Mex67 does interact with mRNAs via different mRNA-binding adaptors. Yra1 (ALY/REF in mammals) is the first adaptor to intervein by facilitating the recruitment of Mex67 to the mRNP (Strasser *et al*., 2000b, Zenklusen *et al*., 2001) while Nab2 has been reported to form a complex with Mex67 and Yra1 (Batisse *et al*., 2009, Iglesias *et al*., 2010). Interestingly, an excess of Nab2 can bypass the loss of Yra1 (Iglesias *et al*., 2010). Npl3, an RNA-binding protein, also directly interacts with Mex67 and participates in mRNA nuclear export (Lee *et al*., 1996).

The assembly of pre-mRNAs into mRNPs is functionally and physically coupled to transcription. Indeed, TREX, a protein complex composed of THO (Hpr1, Tho2, Mft1, and Thp2), the Mex67 adapters Yra1 and the DEAD-box ATPase Sub2 (Strasser *et al*., 2002), is recruited to the transcribing RNAPII and helps in the assembly of an export competent mRNP. THO recruitment to active chromatin is facilitated by the Tho2 subunit (Pena *et al*., 2012). Then, Hpr1 binds Sub2 on nascent transcripts and is implicated in the early recruitment of Mex67, an interaction mediated by a ubiquitin-dependent process (Gwizdek *et al*., 2005, Gwizdek *et al*., 2006, Zenklusen *et al*., 2002). Yra1 is also co-transcriptionally recruited, notably by the cap binding complex (CBC) (Cheng *et al*., 2006, Sen *et al*., 2019, Viphakone *et al*., 2019) and transferred to Sub2 (Johnson *et al*., 2009, Johnson *et al*., 2011). Later, TREX-2 (Sac3, Thp1, Sus1, Cdc31, and Sem1) interacts with the nuclear side of the NPC and participates in the coordination between transcription and mRNA export (Cabal *et al*., 2006, Fischer *et al*., 2004, Jani *et al*., 2009, Rodriguez-Navarro *et al*., 2004).

In contrast to Yra1, which is removed from mRNP complexes before exit the nucleus (Lund *et al*., 2005), Nab2 and Npl3 translocate together with the mRNP and are released at the cytoplasmic face of the NPC (Tran *et al*., 2007). Once the mRNA reaches the cytoplasm, Nup159 and Nup42 NPC subunits recruit two essential mRNA export factors, the DEAD-box ATPase Dbp5 and its ATPase activator Gle1 which are in charge of remodeling and disassembling the mRNPs emerging from the NPC (Adams *et al*., 2017, Hodge *et al*., 2011, Lund *et al*., 2005, Weirich *et al*., 2006). Disassembly of the mRNP in the cytoplasm prevents its return to the nucleus, resulting in unidirectional mRNA translocation. In addition, the release of mRNA export factors allow them to return to the nucleus where they can function in additional rounds of mRNA export (Stewart, 2007). Recently, it was shown that Dbp5 does not affect the interaction of Mex67 with the NPC. Instead, it was proposed that Mex67 does not translocate to the cytoplasmic face of the NPC with the mRNPs but rather functions as a mobile nucleoporin, facilitating their translocation to the cytoplasm (Derrer *et al*., 2019).

Protein ubiquitylation plays very important roles in the control of numerous cellular pathways. In gene expression, ubiquitin has been shown to regulate the activity and turnover of several key transcription factors. The Rsp5 ubiquitin ligase was shown to target RNAPII for degradation during DNA damage (Beaudenon *et al*., 1999, Huibregtse *et al*., 1997, Wu *et al*., 2001). More recently, ubiquitylation of RNAPII in human cells was shown to play a role in transcription-coupled repair (Nakazawa *et al*., 2020, Tufegdzic Vidakovic *et al*., 2020). RNAPII was also shown to be ubiquitylated *in vivo* by the ubiquitin ligase Asr1 (Daulny *et al*., 2008). In this case, the ubiquitylation occurs on transcribed genes and leads to the dissociation of two RNAPII subunits (Rpb4/7), a process involved in the silencing of subtelomeric genes (McCann *et al*., 2016). Ubiquitylation is also involved in mRNA export (Babour *et al*., 2012). The THO component Hpr1 is poly-ubiquitylated by Rsp5 which facilitates the co-transcriptional recruitment of the mRNA export factor Mex67, via its ubiquitin-associated (UBA) domain (Gwizdek *et al*., 2005, Gwizdek *et al*., 2006). The ubiquitin E3 ligase Tom1 is required for ubiquitylation of the Mex67 adaptor Yra1 which promotes its dissociation from mRNP complexes before the export to the cytoplasm (Iglesias *et al*., 2010). H2B ubiquitylation, a histone modification deposited on genes co-transcriptionally, facilitates the assembly of Mex67, Yra1, Nab2 and Npl3 into mRNPs (Vitaliano-Prunier *et al*., 2012). Interestingly, while H2B ubiquitylation mediates the assembly of Npl3 directly, its effect on Mex67, Yra1, Nab2 assembly onto mRNPs involves the ubiquitylation of Swd2, a subunit of the cleavage and polyadenylation factor (CPF), connecting chromatin to transcriptional termination and mRNA export (Vitaliano-Prunier *et al*., 2012). Moreover, a systematic analysis of NPC ubiquitylation was conducted in yeast and showed that more than 50% of Nups can be ubiquitylated (Hayakawa *et al*., 2012). As this modification does not influence Nups localization within the NPC, it is tempting to speculate that it might regulate the interaction with the Mex67 UBA domain during mRNA export.

Here, we identified the deubiquitylase Ubp15 as an RNAPII CTD-interacting protein in *Saccharomyces cerevisiae*. The association of Ubp15 with RNAPII increased in a mutant for the CTD phosphatase Fcp1, suggesting a role in transcriptional elongation. Furthermore, the deletion of *UBP15* rescued the sensitivity of transcription elongation factor mutants to 6-azauracil (6AU), an inhibitor of transcription elongation. While these experiments functionally connect Ubp15 to transcription elongation, the deletion of *UBP15* did not rescue the transcription elongation processivity defect of *dst1Δ* cells in the presence of 6AU, suggesting that the link between Ubp15 and elongation may be indirect. The deletion of *UBP15* rescued the thermosensitivity of a mutant for *RSP5*, an E3 ligase involved in DNA damage and mRNA export. Interestingly, the deletion of *UBP15* suppressed the mRNA export defect of *rsp5* mutants. Finally, we showed that Ubp15 regulates the ubiquitylation of Mex67, which in turn regulates its interaction with THO, a complex involved in mRNA processing and export. Collectively, our data support a role for Ubp15 in coupling transcription to mRNA export.

## RESULTS

### Ubp15 associates with RNAPII and the NPC

To identify new factors interacting with RNAPII in a CTD phosphorylation-dependent manner, we performed a proteomic analysis of RNAPII complexes, affinity-purified from wild type (WT) and *fcp1-1* cells (**Figure 1A, Figure 1–figure supplement 1**). Fcp1 is a major CTD phosphatase and its mutation leads to increased CTD phosphorylation at serines 2, 5 and 7 (Bataille *et al*., 2012). Quantitative analysis of the RNAPII-associated proteins identified 45 proteins differentially associated with the polymerase in *fcp1-1* cells (35 being more abundant and 10 being less abundant by at least twofold; **Figure 1B, Supplementary file 1**). One RNAPII-interacting protein identified in this experiment is Ubp15, a deubiquitylase known for its role in the regulation of endocytosis (Ho *et al*., 2017, Kouranti *et al*., 2010), progression into S phase (Alvarez *et al*., 2016, Ostapenko *et al*., 2015), peroxisomal export (Debelyy *et al*., 2011) and methylmercury susceptibility (Hwang *et al*., 2012), but with no known roles in transcription. The association of Ubp15 with RNAPII is increased in the *fcp1-1* mutant, suggesting it is recruited (directly or indirectly) to the elongating polymerase via CTD phosphorylation. To confirm this interaction, we performed a reciprocal affinity purification experiment where Ubp15 was affinity-purified in WT cells and analyzed by mass spectrometry (**Figure 1C**) and western blot (**Figure 1D**). This experiment confirmed the interaction between Ubp15 and the phosphorylated form of RNAPII (**Figure 1D**) and, surprisingly, revealed the enrichment of almost the entire NPC (**Figure 1C**). Collectively, these experiments identified Ubp15 as an interactor of the phosphorylated RNAPII and the NPC.

**Figure 1.**
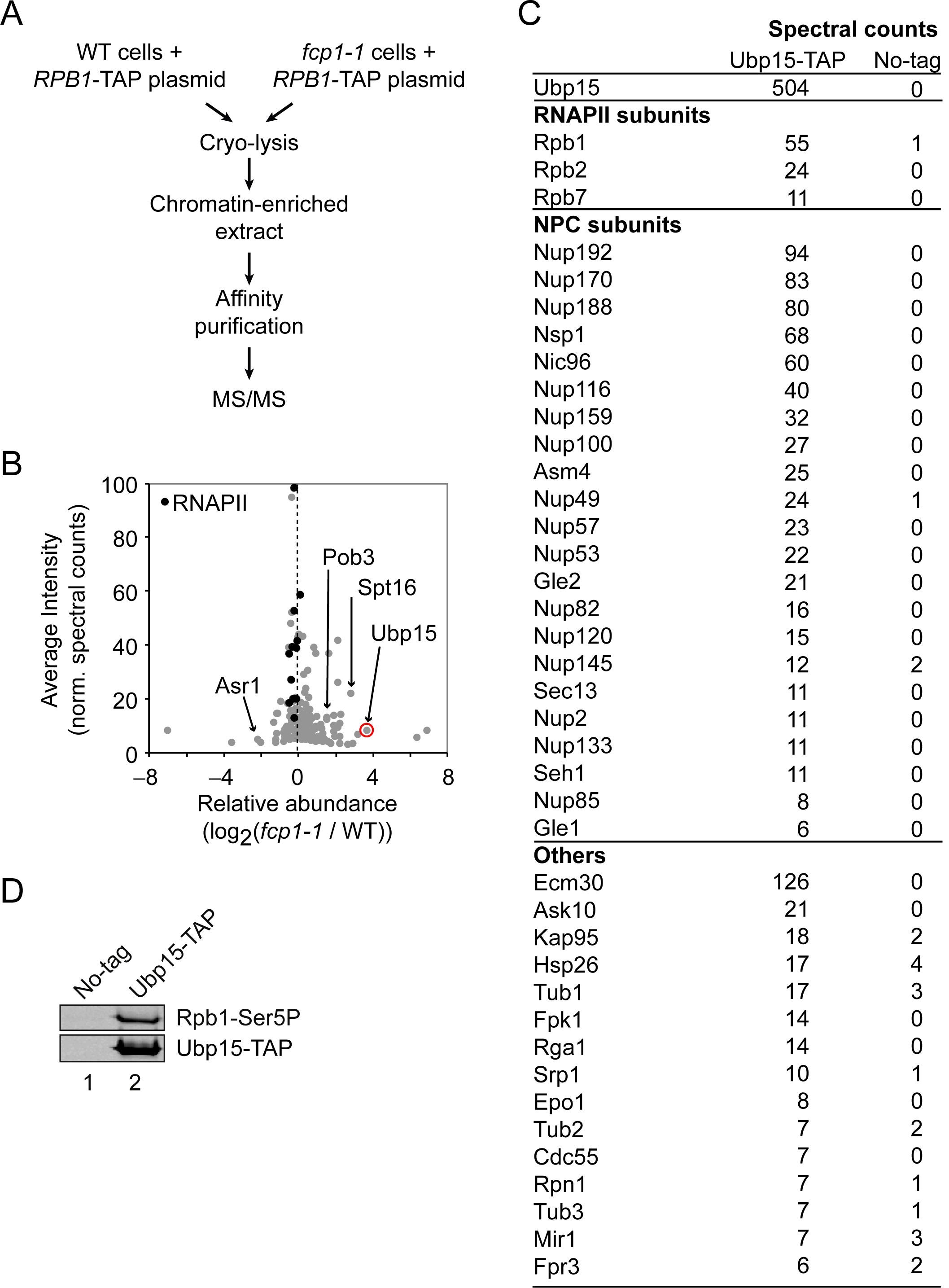
Ubp15 interacts with the phosphorylated RNAPII and with the NPC. **A**) A schematic representation of the proteomic experiment shown in panel B. **B**) Volcano plot showing the average intensity versus relative abundance (log_2_) of proteins identified in RNAPII complexes purified from *fcp1-1* versus WT cells using spectral counts as a proxy for relative protein abundance. RNAPII subunits are shown in black and other proteins of interest are indicated. Ubp15 is circled in red. In the interest of clarity, the maximum value of the y-axis was set at 100, resulting in Rpb1 and Rpb2 not appearing on the graph. See **Supplementary file 1** for the complete list of values. **C**) Spectral counts of the proteins identified in a TAP-tag purification of Ubp15. **D**) Western blot confirming that Ubp15 is associated with a phosphorylated form of RNAPII (Rpb1-Ser5P).

### Ubp15 genetically interacts with elongation factors

Interestingly, the histone chaperone and elongation factor FACT (Spt16 and Pob3) is also enriched in RNAPII purified from *fcp1-1* cells (**Figure 1B, Figure1–figure supplement 1B, Supplementary file 1**) and both FACT and Rpb1 are substrates of Ubp15 in *S*. *pombe* (Beckley *et al*., 2015). In addition, the association of Asr1, a ubiquitin ligase known to bind the CTD and to ubiquitylate RNAPII in the context of elongation (Daulny *et al*., 2008, McCann *et al*., 2016), is decreased in *fcp1-1* cells (**Figure 1B, Supplementary file 1**). These results prompted us to investigate a possible role for Ubp15 in transcription elongation.

First, we looked for sensitivity to 6-azauracil (6AU), an inhibitor of GTP biosynthesis commonly used to test for mutations that affect transcriptional elongation (Riles *et al*., 2004). While the *ubp15Δ* mutant did not show significant sensitivity to 6AU on its own, it rescued the 6AU sensitivity of several elongation factor mutants including *dst1Δ* (TFIIS), *spt4Δ* and *spt5-CTRΔ* (DSIF), *spt6-1004* and *hpr1Δ* (THO) (**Figure 2A, Figure 2–figure supplement 1A**). Interestingly, the 6AU sensitivity of other elongation factors such as *bur2Δ, ctk1Δ, rtf1Δ*, and *cdc73Δ* mutants was not affected by the deletion of *UBP15* (**Figure 2–figure supplement 1A**). Then, we used a Ubp15 catalytic dead mutant, *ubp15*-C214A (Debelyy *et al*., 2011), to test whether the observed genetic interactions were mediated by its catalytic activity. Double mutants *ubp15Δ*/*dst1Δ* and *ubp15Δ/spt4Δ* were complemented with plasmids expressing WT or C214A versions of Ubp15 and spotted on 6AU. The *ubp15*-C214A mutant, but not the WT *UBP15*, rescued the *dst1Δ* and *spt4Δ* sensitivities to 6AU (**Figure 2B**), demonstrating that the effect of Ubp15 on elongation depends on its catalytic activity.

**Figure 2.**
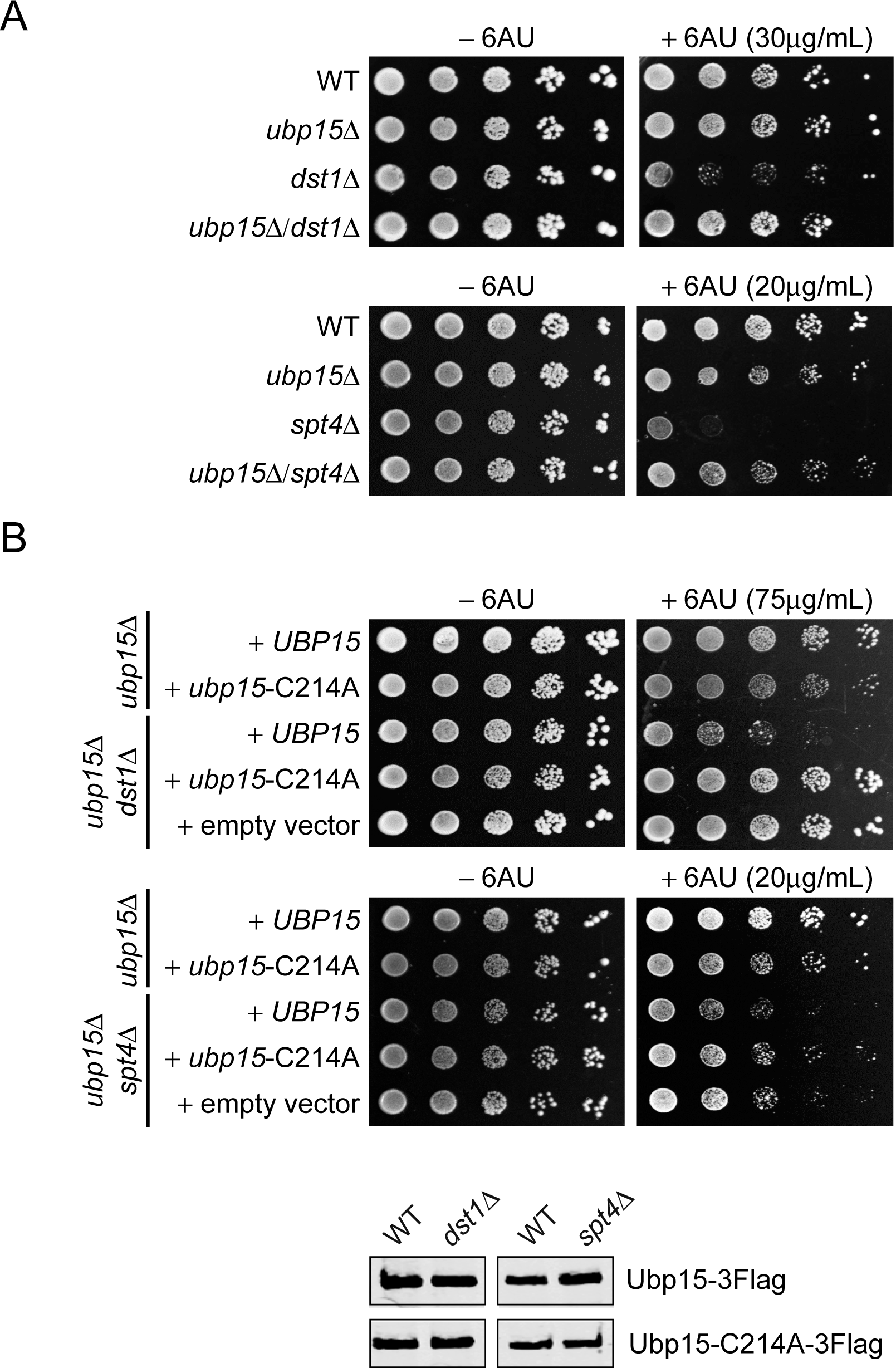
The deletion of *UBP15* suppresses the 6AU sensitivity of *dst1Δ* and *spt4Δ* cells. **A**) Serial-dilution growth assays assessing the 6AU sensitivity of WT, *dst1Δ* and *spt4Δ* cells, alone and in combination with *ubp15*Δ. The indicated yeast strains were grown to saturation in YNB medium lacking uracil (–URA), washed, resuspended at the same density in water, serial diluted (fivefold series) and spotted on –URA in the absence or presence of 6AU as indicated. Plates were incubated at 30°C for 3 days. **B**) Top: Serial-dilution growth assays assessing the requirement of the catalytic activity of Ubp15 for its effect on the 6AU sensitivity of WT, *dst1Δ* and *spt4Δ* cells. Strains were deleted for *UBP15, DST1* and *SPT4*, alone or in combinations, and transformed with empty vector, plasmids expressing Ubp15-3Flag (*UBP15*) or catalytic dead Ubp15-C214A-3Flag (*ubp15*-C214A) and spotted on –URA lacking histidine (–URA/–HIS) in the absence or presence of 6AU as indicated. Bottom: Equal expression levels of WT and catalytic dead versions of Ubp15-3Flag in WT, *dst1Δ* and *spt4Δ* cells were confirmed by Western blot. Note that the 6AU concentration varies and has been optimized for each mutant.

Noteworthy, we also looked at the phenotype of *ubp15Δ* cells under several other growth conditions and confirmed previously described sensitivity to cold temperature and hydroxyurea (HU) (Amerik *et al*., 2000, Ostapenko *et al*., 2015), while elevated temperature, ultraviolet light, formamide or caffeine had no detectable effect (**Figure 2–figure supplement 1B**). Collectively, these genetic interactions confirm previous literature on Ubp15 and demonstrate a functional link between the Ubp15 deubiquitylase activity and transcription elongation.

### Ubp15 does not regulate RNAPII processivity

The suppression of the 6AU sensitivity of several elongation factors is consistent with a role for Ubp15 in transcription elongation. In *S*. *cerevisiae*, elongation factors affect the processivity of RNAPII (that is, the capacity of the polymerase to reach the end of the gene) but have no measurable impact on the elongation rate (Mason *et al*., 2005). We, therefore, tested the effect of *UBP15* deletion on RNAPII processivity in WT and *dst1Δ* cells. We first compared RNAPII occupancy at the beginning and the end of *YLR454W* (a ∼8 kb-long gene) by chromatin immunoprecipitation (ChIP) followed by quantitative PCR (qPCR) as described before (Mason *et al*., 2003, 2005, Schwabish *et al*., 2004, Strasser *et al*., 2002). Surprisingly, we found that the deletion of *UBP15* did not rescue the processivity defect of *dst1Δ* cells in the presence of 6AU (**Figure 3A**). To extend this analysis to the whole genome, we performed ChIP followed by hybridization on tiling microarrays (ChIP-chip) experiments in WT, *ubp15Δ, dst1Δ*, and *ubp15Δ*/*dst1Δ* cells and mapped average RNAPII occupancy over transcribed genes (**Figure 3B, Figure3–figure supplement 1**). These experiments confirmed the processivity defect of *dst1Δ* cells but, consistently with our ChIP-qPCR analysis of the *YLR454W* gene (**Figure 3A**), revealed no effect of *UBP15* deletion on RNAPII processivity in WT or *dst1Δ* cells. This result was somewhat unexpected given the genetic interactions described in **Figure 2** and suggests that *UBP15* deletion may rescue the 6AU sensitivity of elongation factor mutants indirectly. Alternatively, Ubp15 may affect elongation rate, a parameter not investigated here, but this appears unlikely since no elongation factor has been shown to affect this parameter in yeast (Mason *et al*., 2005). One attractive possibility, given the interaction between Ubp15 and the NPC, is that *UBP15* deletion affects a post-transcriptional process in a way that compensates for elongation defects of elongation factor mutants.

**Figure 3.**
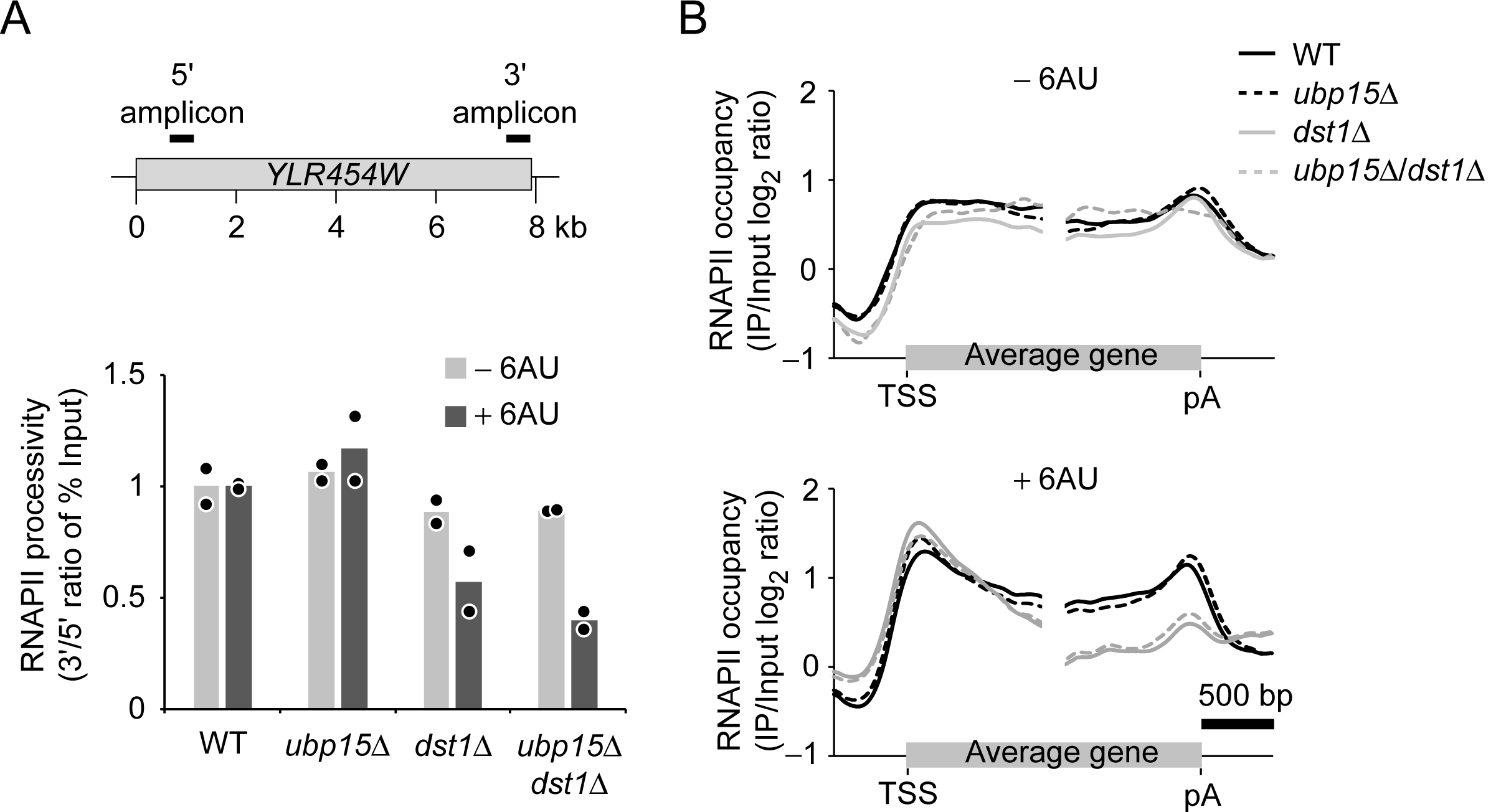
*UBP15* deletion does not rescue RNAPII processivity in *dst1Δ* cells. **A**) RNAPII processivity, defined as the ratio of the % of Input detected in the 3’ amplicon divided by the % of Input detected in the 5’ amplicon, after 30 min treatment with 6AU (dark grey) and absence of 6AU (light grey), as determined by ChIP-qPCR. Experiments were performed in two biological replicates. Bars show the average and circles show individual replicates. The position of PCR amplicons over the *YLR454W* gene used for the qPCR is indicated on the sketch above the graphs. **B**) Aggregate profiles of RNAPII (Rpb3) occupancy over highly expressed yeast genes longer than 1 kb (n=234) as determined by ChIP-chip after 1 hr treatment with 6AU. TSS, transcription start site; pA, polyadenylation site.

### Ubp15 controls nuclear polyA RNA accumulation

To identify a role for Ubp15 that may explain its indirect elongation phenotype, we looked for an E3 ligase that would oppose its function. Through testing several candidates (**Figure 4A, Figure 4–figure supplement 1A**), we noticed that a mutant of the E3 ligase Rsp5 (*rsp5-1*) exacerbated the 6AU sensitivity of *dst1Δ* cells (**Figure 4A**). This effect of *rsp5-1* on the *dst1Δ* 6AU sensitivity is opposite to that of *ubp15Δ* (**Figure 2A**) suggesting that Rsp5 and Ubp15 may oppose each other in regulating a process genetically connected to transcription elongation. Furthermore, the deletion of *UBP15* can partially rescue the 6AU sensitivity of *rsp5-1/dst1Δ* mutant (**Figure 4A**). The antagonistic relationship between Rsp5 and Ubp15 is also supported by the fact that the deletion of *UBP15* rescued the thermosensitivity phenotype of *rsp5-ts* mutants (**Figure 4B** and **Figure 4–figure supplement 1B**). Importantly, the rescue of *rsp5-1* thermosensitivity was also observed with the *ubp15-C214A* mutant indicating it is dependent on the catalytic activity of Ubp15 (**Figure 4C**).

Rsp5 is an E3 ligase involved in DNA damage (Beaudenon *et al*., 1999) and mRNA export (Rodriguez *et al*., 2003). At high temperatures, the *rsp5-1* mutant accumulates mRNA in the nucleus (Rodriguez *et al*., 2003). We, therefore, compared polyA RNA localization in *rsp5-1* and *ubp15Δ*/*rsp5-1* cells by RNA fluorescent in situ hybridization (FISH) using a Cy5-labelled oligo-dT_45_ probe. Interestingly, these experiments showed that the deletion of *UBP15* suppressed the mRNA export defect of the *rsp5-1* mutant (**Figure 4D**). Indeed, the deletion of *UBP15* reduced the number of cells retaining polyA RNAs in their nucleus from around 75% (172/222) to less than 5% (11/227). This suppression of mRNA export defect likely explains how *UBP15* deletion restored viability of *rsp5* mutants and clearly establishes a role for Ubp15 in mRNA export.

**Figure 4.**
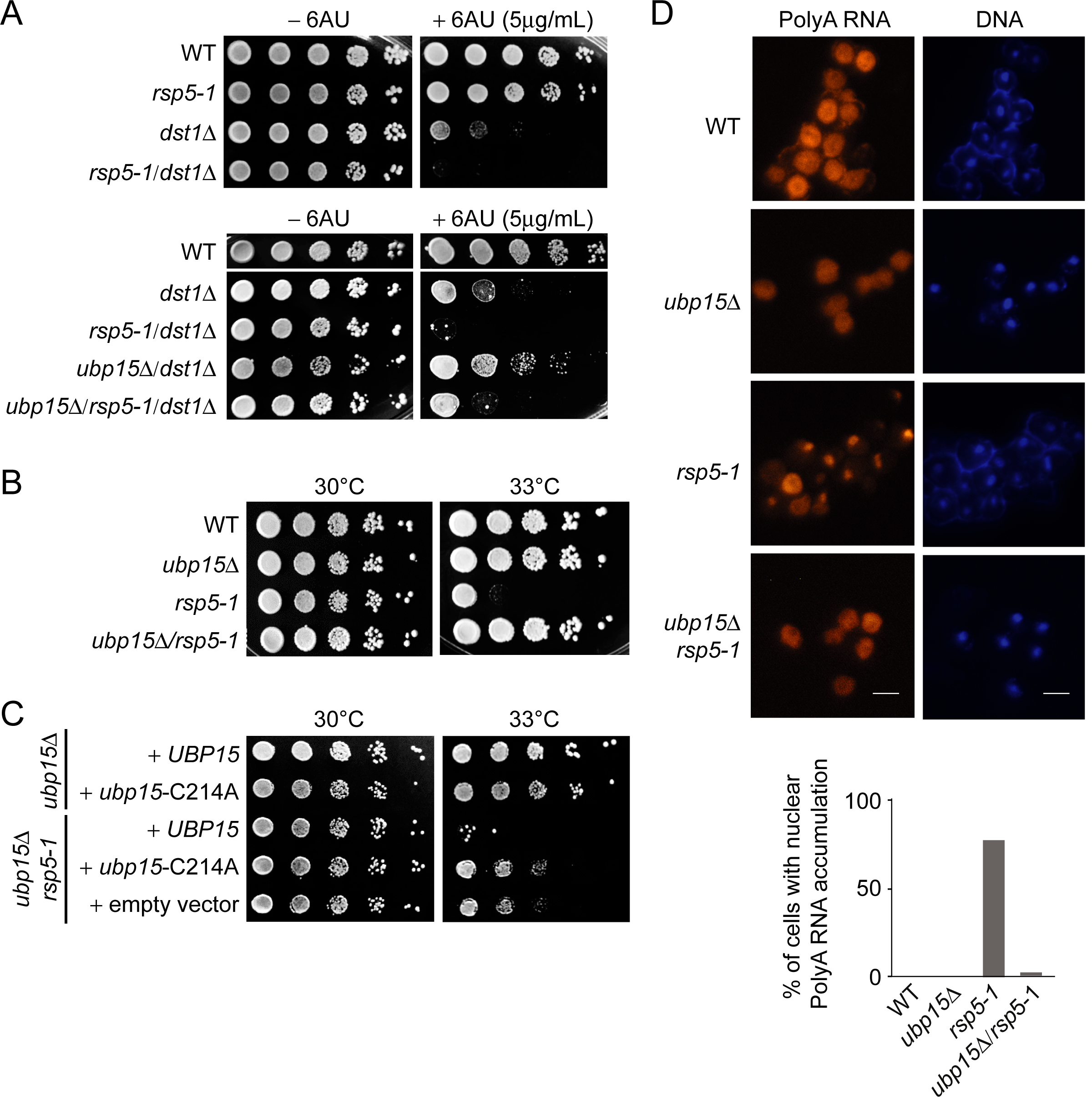
Deletion of *UBP15* rescues phenotypes of *rsp5-1* mutants. **A**) Serial-dilution growth assays assessing the sensitivity of *rsp5-1* cells, alone or in combination with *dst1*Δ and or *ubp15*Δ, to 6AU. The indicated yeast strains were grown to saturation in YNB medium lacking uracil (–URA), washed, resuspended at the same density in water, serial diluted (fivefold series) and spotted on –URA in the absence or presence of 6AU as indicated. Plates were incubated at 30°C for 3 days. **B**) Serial-dilution growth assays assessing the effect of *UBP15* deletion on the viability of *rsp5-1* cells at 33°C. The indicated yeast strains were grown to saturation in YPD, washed, resuspended at the same density in water, serial diluted (fivefold series) and spotted on YPD. Plates were incubated for 3 days at 30°C or 33°C as indicated. **C**) Serial-dilution growth assays assessing the contribution of the catalytic activity of Ubp15 to the genetic interaction between *RSP5* and *UBP15* shown in panel B. Strains deleted for *UBP15*, alone or in combination with the *rsp5-1* mutation, were transformed with empty vector, plasmids expressing Ubp15-3Flag (*UBP15*) or catalytic dead Ubp15-C214A-3Flag (*ubp15*-C214A), spotted on YNB medium lacking histidine (–HIS) and incubated at 30°C or 33°C as indicated. **D**) RNA FISH experiments looking at bulk polyA RNAs in WT, *ubp15Δ, rsp5-1* and *ubp15Δ*/*rsp5-1* cells (FY genetic background). The indicated strains were grown at 30°C in YPD then shifted to 37°C for 3 hr before being analyzed by FISH using Cy5-oligo-dT_45_. DNA was stained with DAPI. Scale bar, 10 µm. The percentage of cells (from at least 200 cells in each strain) with retention of polyA RNA in the nucleus is indicated on the graphic shown at the bottom (WT: 0/200, *ubp15Δ:* 0/200, *rsp5-1*: 172/222, *ubp15Δ*/*rsp5-1*: 11/227).

### Ubp15 and Rsp5 have opposite effects on Mex67 ubiquitylation

To identify potential substrates for Ubp15 in the mRNA export pathway, we measured ubiquitylation levels of Mex67 (**Figure 5B,C**), six NPC components (Nup82, Nup133, Nup57, Nup120, Nup145, and Nup159) (**Figure 5–figure supplement 1A**) and four nuclear export factors (Hpr1, Nab2, Npl3, and Mtr2) (**Figure 5–figure supplement 1B**) in *ubp15Δ* cells using a previously described *in vivo* ubiquitylation assay (**Figure 5A)** (Gwizdek *et al*., 2005). These experiments identified Mex67 as a likely substrate of Ubp15 (**Figure 5B,C**). Interestingly, Mex67 ubiquitylation levels decreased in *rsp5-1* (**Figures 5B**), suggesting that Rsp5 and Ubp15 may oppose each other in the control of Mex67 ubiquitylation. To assess this possibility more directly, we tested Mex67 ubiquitylation levels in *ubp15Δ*/*rsp5-1* double mutant and found that the deletion of *UBP15* normalized Mex67 ubiquitylation of *rsp5-1* cells (**Figure 5C**). Collectively, these experiments established Rsp5 and Ubp15 as an E3 ligase/deubiquitylase tandem regulating Mex67 ubiquitylation.

**Figure 5.**
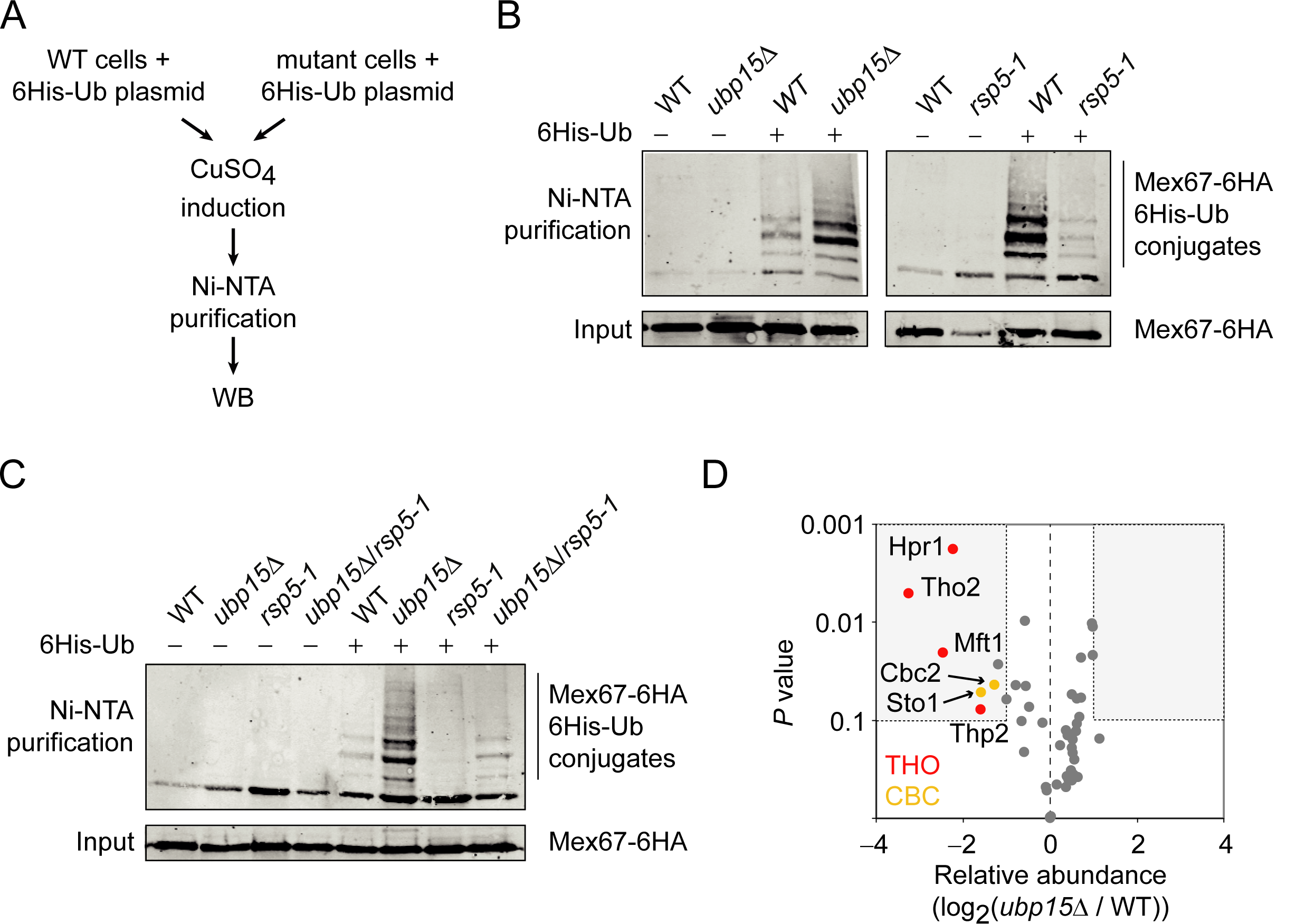
The ubiquitylation of Mex67 is regulated by Ubp15 and Rsp5. **A**) A schematic representation of the *in vivo* ubiquitylation assay used in panels B and C. A plasmid expressing polyhistidine-tagged ubiquitin (6His-Ub) under the control of a copper-inducible promoter was transformed in WT and mutant cells. 6His-Ub expression was induced with copper sulfate (CuSO_4_) and His-tagged ubiquitin-conjugated proteins were purified using Ni-NTA beads and analyzed by Western Blot. **B**) Western blots for Mex67-6HA levels from His-tagged ubiquitin-conjugated protein pulldowns (Ni-NTA) and their inputs expressing (+) or not (–) 6His-Ub in WT, *ubp15Δ*, and *rsp5-1* cells. **C**) Same as panel B but for WT, *ubp15Δ, rsp5-1* and *ubp15Δ*/*rsp5-1* cells. **D**) Ubp15 regulates the interaction of THO and CBC with Mex67. A volcano plot showing the significance versus the log_2_ fold change for the proteins identified in Mex67-3Flag purifications by MS in *ubp15Δ* versus WT cells. Grey dots show the bulk of the data and the regions for significant (*P* value<0.1) twofold changes are boxed. THO and CBC subunits are labeled in red and gold, respectively. See **Supplementary file 2** for the complete list of values.

### Ubp15 regulates the interaction of Mex67 with THO and the cap-binding complex

Our data so far show that Rsp5 and Ubp15 oppose each other in the regulation of mRNA export and ubiquitylation of Mex67. Since Mex67 is known to function as part of a large and dynamic protein interaction network, we hypothesized that its ubiquitylation may modulate its interactome. Hence, we tested whether the proteome associated with Mex67 is modified in *ubp15Δ* cells. We performed a proteomic analysis of Mex67, affinity-purified from WT and *ubp15Δ* cells (**Figure 5D, Figure 5–figure supplement 2**). As expected, based on previous studies (Batisse *et al*., 2009, Gwizdek *et al*., 2006, Oeffinger *et al*., 2007, Santos-Rosa *et al*., 1998, Saroufim *et al*., 2015, Strasser *et al*., 2000b, Zenklusen *et al*., 2001), we found a large number of Mex67-interacting proteins including Mtr2, Nab2, Yra2, the THO complex and nearly the entire NPC (**Supplementary file 2**). Among those, only seven were differentially associated with Mex67 in *ubp15Δ* cells by at least twofold (**Figure 5D**). Interestingly, all those interactors were less associated with Mex67 in *ubp15Δ* cells, suggesting that the ubiquitylation of Mex67 negatively regulates its association with these factors. Strikingly, these proteins include all four subunits of the THO complex (Hpr1, Tho2, Mft1, and Thp2) and both subunits of the heterodimeric cap-binding complex (CBC; Sto1 and Cbc2) known to recruit THO/TREX to the 5’-cap of nascent pre-mRNAs (**Figure 5D**) in yeast and humans (Cheng *et al*., 2006, Sen *et al*., 2019, Viphakone *et al*., 2019). Together, these results support a model where Mex67 ubiquitylation, controlled by Rsp5 and Ubp15, regulates the association of Mex67 to pre-mRNAs via interactions with THO and cap binding factors.

## DISCUSSION

In this study, we characterized the role of the deubiquitylase Ubp15 in mRNA export in yeast. Initially, we identified Ubp15 as an interactor of the phosphorylated RNAPII and the NPC. We then investigated for a role in transcription elongation and showed that the deletion of *UBP15* rescues the sensitivity of diverse transcription elongation factor mutants to 6AU but surprisingly does not rescue the transcription elongation processivity defect in these mutants. These results argue against a direct role of Ubp15 in transcriptional elongation and rather suggest that it regulates a post-transcriptional event in a way that indirectly suppresses the growth phenotype of elongation factor mutants. While looking for an E3 ligase that may oppose the function of Ubp15, we found that the deletion of *UBP15* can rescue the thermosensitivity of a mutant of *RSP5*. Interestingly, the deletion of *UBP15* suppresses the mRNA export defect of *rsp5* ts mutants, establishing a role for Ubp15 in mRNA export. We then showed that Ubp15 and Rsp5 control the ubiquitylation levels of Mex67 and that Mex67 ubiquitylation negatively regulates its interaction with THO and CBC. Collectively, our data support a role for Ubp15 in coupling transcription to mRNA export.

Our initial data showed that the interaction of Ubp15 with RNAPII is increased in the mutant for the CTD phosphatase *fcp1-1*. In this mutant, RNAPII is hyper-phosphorylated, notably on its Ser2, a phosphorylation state which is predominant towards the 3’-end of genes (Bataille *et al*., 2012). mRNA processing and export factors also function near the 3’-end of genes and some even require Ser2 phosphorylation for their recruitment (Jeronimo *et al*., 2013), which could explain the link between Ubp15, RNAPII CTD phosphorylation, and mRNA export.

In principle, one possible role of Mex67 ubiquitylation/deubiquitylation by Rsp5 and Ubp15, respectively, could be to regulate its nuclear localization. We, however, could not observe any effect of *ubp15Δ* or *rsp5-1* mutants on Mex67 localization (data not shown). This is consistent with a recent paper from Derrer et al. which demonstrated that Mex67 localization is restricted to the NPC (Derrer *et al*., 2019). Instead, we showed that Mex67 ubiquitylation negatively affects its interaction with the THO and CBC complexes. Interestingly, CBC was shown to recruit THO/TREX to the 5’-cap via an interaction with Yra1 (Cheng *et al*., 2006, Sen *et al*., 2019, Viphakone *et al*., 2019). Hence, we propose that the assembly of Mex67 into mRNPs during transcription is counteracted by its ubiquitylation. In such a model, Ubp15, recruited by the phosphorylated RNAPII CTD, would antagonize Mex67 ubiquitylation, hence allowing Mex67 recruitment to the pre-mRNA by CBC and THO/TREX. Ubiquitylation of Mex67 may prevent interaction with CBC and THO/TREX by a simple sterical block or, –since Mex67 contains the UBA domain– Mex67 ubiquitylation may trigger an intramolecular interaction between the conjugated ubiquitin and the UBA domain, creating a folded closed conformation. In line with such a scenario, the UBA is known to bind ubiquitylated Hpr1 (a component of THO/TREX) (Gwizdek *et al*., 2006). A competition of the UBA domain for binding to ubiquitylated Hpr1 and the internal Mex67 ubiquitylated site would provide a powerful switch for Mex67 binding to THO/TREX.

Interestingly, Hpr1 ubiquitylation is mediated by Rsp5 (Gwizdek *et al*., 2005, Gwizdek *et al*., 2006). Hence, Rsp5-mediated ubiquitylation of Hpr1 and Mex67 deubiquitylation by Ubp15 would work hand in hand towards the assembly of Mex67 into mRNPs. This model, however, is at odds with our observation that *UBP15* deletion and catalytically inactive Ubp15 do revert Rsp5 export defects. This conundrum may be solved by considering that Mex67 and Hpr1 ubiquitylation need to turnover for mRNA export to optimally function. Indeed, it sounds reasonable to think that Mex67 and Hpr1 need to flip back and forth between their ubiquitylated and non-ubiquitylated forms considering the very dynamic nature of the interactions that they form and the fact that mRNPs are being remodeled during their journey from the transcription site to the cytoplasmic side of the nuclear envelope. Alternatively, in *rsp5* mutant cells, Mex67 –which lost the ability to interact with THO/TREX via Hpr1– may (hypothetically) find another route towards assembly into mRNPs when ubiquitylated, a condition that would be favored by the deletion of *UBP15*. This may be reminiscent of what happens during heat-shock, where stress-response mRNAs are rapidly exported by Mex67, without the need for adapters (Zander *et al*., 2016). The exact molecular mechanisms describing the role of Ubp15 in mRNA export and its coupling to transcription will require additional work, but the data presented here clearly establish this deubiquitylase as a new player in this arena.

## MATERIALS AND METHODS

### Yeast strains and plasmids

Genotypes for the yeast strains used in this study are listed in **Supplementary file 3**. All tagged and deletion strains were done by homologous recombination of appropriate PCR cassettes. The catalytic dead *ubp15* mutation (C214A) was introduced into pFR559 by inverted PCR (forward: p-GCCTATTTGAATTCGTTATTGC, reverse: TGTGGCACCCTGATTTCGGAAGCCA) to generate pFR560 (see **Supplementary file 4**). Each construct was validated by sequencing and their expression tested by western blot analysis.

### Serial-dilution growth assays

Cells were grown to saturation in the indicated media at 30°C, washed and resuspended to an OD_600_ of 1.0 in H_2_O. Cells were then subjected to fivefold serial dilutions and spotted onto the appropriate media. Plates were incubated at 30°C for 3 days unless specified otherwise. Images presented in figures are representative examples of at least two biological replicates, except for the cold sensitivity (15°C) assay in **Figure 2-figure supplement 1B** and thermosensitivity assays for r*sp5-sm1* and *rsp5-3* in **Figure 4-figure supplement 1B**, which were done once. In addition, several phenotypes were confirmed in two different yeast genetic backgrounds.

### Purification of proteins associated with RNAPII, Ubp15, and Mex67

TAP-tagged Rpb1 subunit of RNAPII from WT and *fcp1-1* cells was purified in two biological replicates by one-step affinity purification essentially as published previously (Jeronimo *et al*., 2015). In brief, following cryogenic disruption of cells (Trahan *et al*., 2016), frozen cell grindate (5 g) was thawed into nine volumes of EB150 extraction buffer (20 mM Tris-HCl pH 7.5, 150 mM KOAc, 1 mM EDTA pH 8.0, 0.5% Triton X-100, 10% glycerol, 1 mM DTT, protease and phosphatase inhibitors and 1:5000 Antifoam B (Sigma)), vortexed for 30 sec and homogenized (Polytron PT 1200E; Kinematica AG) for another 30 sec, to allow for maximal recovery of chromatin proteins. The cleared extract was incubated for 1 hr at 4°C with 200 µL of pre-washed magnetic Dynabeads M-270 Epoxy (Thermo Fisher Scientific) conjugated to rabbit IgG (Sigma). Dynabeads were then collected and washed five times with EB150 buffer and two times with TEV150 protease cleavage buffer (10 mM Tris-HCl pH 8.0, 150 mM KOAc, 0.5 mM EDTA pH 8.0, 0.1% Triton X-100, 10% glycerol and 1mM DTT). The isolated protein complex was eluted by incubating the beads overnight at 4°C with 200 units of TEV protease (Thermo Fisher Scientific) in 500 µl of TEV150 buffer. After digestion, the collected eluate was incubated with pre-washed nickel-nitrilotriacetic acid (Ni-NTA) agarose beads (Qiagen) for 90 min at 4°C to remove the His-tagged TEV protease. The collected eluate was concentrated to ∼300 µl by dialysis in PEG dialysis buffer (10 mM HEPES-KOH pH 7.9, 0.1 mM EDTA pH 8.0, 100 mM KOAc, 20% glycerol, 20% PEG-8000 and 1 mM DTT) and then dialyzed in No-PEG dialysis buffer (10 mM HEPES-KOH pH 7.9, 0.1 mM EDTA pH 8.0, 100 mM KOAc, 20% glycerol and 1 mM DTT). A fraction of the purified proteins was separated by SDS-PAGE on a 4-12% NuPAGE Novex Bis-Tris precast gel (Thermo Fisher Scientific) and visualized by silver staining and western blot analysis with the indicated antibodies. Proteins were precipitated with TCA before mass spectrometry analysis.

For the purification of the Ubp15-TAP complex, the same one-step affinity purification procedure was used except that the salt concentration of the buffers involved in the preparation of the extract and the purification step was increased to 500 mM. In brief, EB500 extraction buffer (20 mM Tris-HCl pH 7.5, 500 mM KOAc, 1 mM EDTA pH 8.0, 0.5% Triton X-100, 10% glycerol, 1 mM DTT, protease and phosphatase inhibitors and 1:5000 Antifoam B (Sigma)) and TEV500 protease cleavage buffer (10 mM Tris-HCl pH 8.0, 500 mM KOAc, 0.5 mM EDTA pH 8.0, 0.1% Triton X-100, 10% glycerol and 1mM DTT) were used.

For the purification of the proteins associated with Mex67-3Flag from WT and *ubp15Δ* cells, frozen cell grindate (1 g) was thawed into nine volumes of TBT extraction buffer (20 mM HEPES-KOH pH 7.5, 110 mM KOAc, 1 mM MgCl_2_, 0.5% Triton X-100, 0.1% Tween 20, 1 mM DTT, protease inhibitor mixture, SUPERaseIn RNase inhibitor (Thermo Fisher Scientific) and 1:5000 Antifoam B (Sigma)) as previously described (Oeffinger *et al*., 2007). The cell extracts were then vortexed for 30 sec, homogenized (Polytron PT 1200E; Kinematica AG) for another 30 sec and clarified by centrifugation at 3,500 rpm for 10 min at 4°C. The Mex67-3Flag tagged protein was isolated using 200 µL of pre-washed magnetic Pan Mouse Dynabeads (Thermo Fisher Scientific) coupled to 34 µg of anti-Flag M2 mouse monoclonal antibody (Sigma). After binding for 1 hr at 4°C, Dynabeads were collected and washed five times with TBT extraction buffer and five times with TBT washing buffer (20 mM HEPES-KOH pH 7.5, 110 mM KOAc, 1 mM MgCl_2_). The isolated protein complex was eluted twice by incubating each time the beads 20 min at room temperature with 500 µl of NH_4_OH elution buffer (0.5 M NH_4_OH, 1 mM EDTA pH 8.0). The pooled eluates were splitted into four aliquots, then dried in a speed-vac at room temperature. One aliquot of purified proteins was separated by SDS-PAGE on a 4-12% NuPAGE Novex Bis-Tris precast gel and visualized by silver staining. Another aliquot was analysed by mass spectrometry.

### Protein identification by mass spectrometry

The experiments were essentially performed as described previously (Bataille et al. 2012). Protein samples were re-solubilized in 6 M urea buffer followed by reduction and alkylation before digestion with trypsin (Promega) at 37°C for 18 hr. The digested peptide mixtures were dried down in vacuum centrifuge and stored at −20°C until LC-MS/MS analysis. Prior to LC-MS/MS, the digested peptide mixtures were resolubilized in 0.2% formic acid and desalted/cleaned up by using C18 ZipTip pipette tips according to the manufacturer’s instructions (Millipore). Eluates were dried down in vacuum centrifuge and then re-solubilized in 2% ACN / 1% formic acid. The LC column used was a C18 reversed-phase column packed with a high-pressure packing cell. A 75 µm i.d. Self-Pack PicoFrit fused silica capillary column of 15 cm long (New Objective) was packed with the C18 Jupiter 5 µm 300Å reverse-phase material (Phenomenex). This column was installed on the Easy-nLC II system (Proxeon Biosystems) and coupled to the LTQ Orbitrap Velos or the Orbitrap Fusion (Thermo Fisher Scientific) equipped with a Proxeon nanoelectrospray ion source.

### Mass spectrometry data analysis

Protein database searching was performed with Mascot 2.2 or 2.6 (Matrix Science) against the *S*. *cerevisiae* NCBInr protein database (2010-12-14 release) or the UniProt_Saccharomyces_Cerevisiae (559292 - strain ATCC 204508 / S288c) database. The mass tolerances for precursor and fragment ions were set to 15 ppm and 0.60 Da, respectively. Trypsin was used as the enzyme allowing for up to two missed cleavages. Carbamidomethyl and oxidation of methionine were allowed as variable modifications. Data interpretation was performed using Scaffold 3.1.2 or 4.8 (Proteome Software). Spectral counts values were exported in Excel and processed as follows.

For RNAPII purifications in WT and *fcp1-1* cells, spectral counts from two biological replicates of Rpb1-TAP purified from WT cells and two biological replicates of Rpb1-TAP purified from *fcp1-1* cells, together with a collection of no-tag controls, were used as a measure of protein abundance. Spectral counts for the 706 identified proteins were floored to 0.1 and normalized to the bait protein level (Rpb1). Proteins with an average spectral count in no-tag controls above 10 (n=44) were removed. Proteins with less than five average spectral counts in both WT and *fcp1-1* conditions (n=495) were also removed. The final dataset contains 170 proteins. For these, the log_2_ relative abundance (log_2_ (*fcp1-1*/WT)) and the average intensity (across all four samples) were calculated. These values are available in **Supplementary file 1** and displayed in **Figure 1B**.

For the Ubp15-TAP purification, spectral counts from Ubp15-TAP and a no-tag control were used as a measure of protein abundance. Proteins with more than 10 spectral counts in no-tag control (n=91) were removed. A protein with less than twofold more spectral count in the tag versus no-tag sample was also removed. Proteins with less than five spectral counts in the Ubp15-TAP sample (n=126) were also removed. Finally, duplicated proteins (different IDs referring to the same protein) were removed (n=3). This analysis led to a final dataset of 41 proteins displayed in **Figure 1C**.

For Mex67-3Flag purifications in WT and *ubp15Δ* cells, spectral counts from three biological replicates of Mex67-3Flag from WT, *ubp15Δ* and a no-tag control were used as a measure of protein abundance. Proteins with less than fivefold enrichment in tagged versus no-tag, and with an average of less than eight spectral counts in tagged experiments were removed. This analysis resulted in 47 proteins displayed in **Figure 5D**. The data were normalized to set the ratio of the spectral count of the bait (Mex67) to 1 between the WT and *ubp15Δ* samples. The log_2_ of the ratio between the normalized average spectral count in *ubp15Δ* and WT cells were computed and a T-test on the spectral counts of the three WT and three *ubp15Δ* samples was used to assess significance.

### ChIP-qPCR

To assess RNAPII (Rpb3) binding along the *YLR454W* gene, ChIP experiments were performed in two biological replicates as previously described (Collin *et al*., 2019). In brief, yeast cells containing the *GAL1* promoter controlling the *YLR454W* gene were inoculated from an overnight preculture in 100 mL of yeast nitrogen-based (YNB) medium lacking uracil (–URA) supplemented with 2% galactose and 2% raffinose (to induce *YLR454W* gene expression). Cells were grown at 30°C until OD_600_ reaches 0.6–0.8 and then 50 mL were treated with 100 µg/mL 6AU for 30 min while the rest was left untreated. Cells were cross-linked with 1% formaldehyde for 30 min at room temperature and quenched with 125 mM glycine. Immunoprecipitation of RNAPII was done using 3 μL of Rpb3 antibody (W0012 from Neoclone) coupled to magnetic Pan Mouse Dynabeads (Thermo Fisher Scientific). ChIP DNA was analyzed by qPCR using primers targeting the 5’ (forward: ACGCAAAGGAACTAGAGAACG, reverse: AATAGGACTCTCCGCCTTGTT) and 3’ (forward: GGTCACAGATCTATTACTTGCCC, reverse: TTCAGGCTCCGTGTAGGAATTA) regions of the *YLR454W* open reading frame. 1% of each Input sample was analyzed in parallel and enrichments were expressed as percent of Input using the following formula: 100*2^[Ct_Input_-6.644-Ct_IP_].

### ChIP-chip

ChIP-chip experiments from WT, *ubp15Δ, dst1Δ* and *ubp15Δ*/*dst1Δ* cells were performed in two biological replicates (except for the *ubp15Δ*/*dst1Δ* strain in the absence of 6AU which was done only once) as previously described (Collin *et al*., 2019). In brief, yeast cells were inoculated from an overnight preculture in 50 mL of YNB–URA supplemented with 2% glucose. Cells were grown at 30°C until OD_600_ reaches 0.6–0.8 and treated (or not) with 100 µg/ml 6AU for 60 min. Cells were cross-linked with 1% formaldehyde for 30 min at room temperature and quenched with 125 mM glycine. Immunoprecipitation of RNAPII was done using 3 μL of Rpb3 antibody (W0012 from Neoclone) coupled to magnetic Pan Mouse Dynabeads (Thermo Fisher Scientific). ChIP and Input samples were labeled with mono-reactive NHS ester fluorescent Cy5 and Cy3 dyes (GE Healthcare) respectively, combined and hybridized for at least 18 hr on custom-designed Agilent microarrays containing 180,000 Tm-adjusted 60-mer probes covering the entire yeast genome with virtually no gaps between probes (Jeronimo *et al*., 2014). Microarrays were washed and scanned using an InnoScan900 (Innopsys) at 2 μm resolution.

### ChIP-chip data analysis

The ChIP-chip data were normalized using the Limma Loess method and replicates were combined as described previously (Ren *et al*., 2000). The data were subjected to one round of smoothing using a Gaussian sliding window with a standard deviation of 100 bp to generate data points in 10 bp intervals (Guillemette *et al*., 2005). Aggregate profiles were generated using the Versatile Aggregate Profiler (VAP) (Brunelle *et al*., 2015, Coulombe *et al*., 2014). In these analyses, only genes that are at least 1 kb long and with an average RNAPII enrichment log_2_ ratio over 1 were retained. Genes were virtually split in the middle and their 5’ and 3’ halves were aligned on the TSS and polyA site respectively. The signal was then averaged in 10 bp bins. Coordinates of TSS and polyA sites are from (Xu *et al*., 2009). Violin plots of RNAPII processivity were built by calculating the log_2_ ratio of Rpb3 occupancy in the last versus first 300 bp for each gene using VAP. The plots were generated using PlotsOfData (Postma *et al*., 2019).

### RNA FISH

RNA FISH was performed as described previously (Babour *et al*., 2016) with a few modifications. Briefly, cells were grown in 50 mL of YPD medium to an OD_600_ of 0.6-0.8 and fixed with 4% paraformaldehyde for 45 min at room temperature on a rotating wheel. Cells were washed twice with cold phosphate buffer (100 mM KHPO_4_ pH 6.4) and once with cold spheroplast buffer (phosphate buffer supplemented with 1.2 M sorbitol). Digestion of yeast cell wall was performed at 30°C with 250 µg of zymolyase 100T (US Biological). Spheroplasts were carefully washed twice and resuspended in 1 mL of cold spheroplast buffer and 200 µL were attached to poly-L-lysine (Sigma)-coated coverslips for 30 min at room temperature. Unadhered cells were washed off and coverslips were stored in 70% ethanol at –20°C for at least 2 hr. Hybridization was carried out in hybridization buffer (50% formamide, 10% dextran sulfate, 4X SSC, 1X Denhardts, 125 μg/ml E. coli tRNA, 500 μg/ml ssDNA, 10 mM ribonucleoside-vanadyl complex (NEB)) supplemented with 50 ng of Cy5-oligo-dT_45_ probe for 12 hr at 37°C in the dark. Coverslips were washed twice with 2X SSC at 37°C for 15 min, once with 1X SSC at room temperature for 15 min and twice with 0.5X SSC at room temperature for 15 min and finally with 1X PBS buffer containing 0.5 µg/mL 4′,6-diamidino-2-phenylindole (DAPI) and mounted onto ProLong Gold antifade reagent (Thermo Fisher Scientific) mounting media.

### Fluorescence microscopy

FISH images were acquired with a Retiga EXi aqua camera (mount on 0.70X) mounted on a DM5500B upright microscope (Leica) and magnified through a 100X oil immersion objective (NA = 1.3). Images of Fluorescent probes were excited using an X-Cite Series 120Q light source (Lumen Dynamics) with the appropriate filters. All hardware parts were controlled with Volocity v5.0 software. Data presented in figures are representative fields of images from two biological replicates. For calculating the percentage of mRNA accumulation in the different strains, we manually visualized 150 to 300 cells and counted the ones with visible polyA RNA accumulation in the nucleus.

### In vivo ubiquitylation assay

Ubiquitylated proteins were detected essentially as described previously (Muratani *et al*., 2005) with some modifications. Yeast strains transformed with a plasmid expressing polyhistidine-tagged ubiquitin (6His-Ub) under the control of a copper-inducible promoter (pFR453) or its non-tagged control (pFR452) (see **Supplementary file 4**) were grown in 50 mL of YNB–URA. When OD_600_ reached 0.6-0.8, cells were induced with 500 µM CuSO_4_ for 2-3 hr at 30°C. Cell cultures were centrifuged and washed twice with ice-cold water, flash-frozen and stored at –80°C. Cell pellets were resuspended in 1 mL of freshly prepared A2 buffer (6 M guanidine-HCl, 100 mM Na_2_HPO_4_/NaH_2_PO_4_ pH 8.0, 10 mM imidazole, 250 mM NaCl, 0.5% NP40) and lysed by glass bead beating 5 min twice. Cell Lysates were then clarified by centrifuging for 15 min at max speed at 4°C. 10 µL of cleared extract were kept as Input and the rest was used for Ni-NTA purification. His-tagged ubiquitin-conjugated proteins were purified by adding 100 µL of 50 % Ni-NTA agarose beads (Qiagen) equilibrated in A2 buffer to 1 mL of cleared extracts and incubated with rotation for 2-4 hr at room temperature. Beads were pelleted and washed twice with 1 mL of A2 buffer, twice with 1 mL of A2/T2 buffer (1 volume A2 buffer + 3 volumes T2 buffer (50 mM Na_2_HPO_4_/NaH_2_PO_4_ pH 8.0, 20 mM imidazole, 250 mM NaCl, 0.5% NP40)) and twice with 1 mL of T2 buffer. Samples were rotated 5 min at room temperature for each wash. After the final wash, all liquid was removed from beads and beads were resuspended in 50 µL of 2X Laemmli buffer supplemented with 250 mM imidazole. Inputs were prepared as previously described (Pepinsky, 1991). Briefly, 800 µL of 100% ethanol was added to 10 µL of cleared extract, vortexed and incubated for 1 hr at –20°C. Inputs were then centrifugated and washed twice with 100% ethanol for 15 min at –20°C. Ethanol was removed and, when completely dry, pellets were resuspended in 100 µL of 2X Laemmli buffer. All samples were boiled for 5 min, centrifuged and supernatants were run on SDS-PAGE for western blot analysis with anti-HA F7 mouse monoclonal antibody (Santa Cruz Biotechnology). Images presented in figures are representative examples of at least three (**Figure 5**) or two (**Figure 5-figure supplement 1**) biological replicates, except for Nup133, which was assayed only once.

## Acknowledgments

This work was funded by grants from the Natural Sciences and Engineering Research Council of Canada (NSERC) to F.R. and the Canadian Institutes of Health Research (CIHR) to F.R (MOP-162334) and J.C. (FDN-143314). F.R. holds a Research Chair from the « Fonds de Recherche Québec –Santé » (FRQS). J.C. holds the Canada Research Chair in Chromatin Biology and Molecular Epigenetics. F.E. held fellowships from « Fondation pour la Recherche Médicale » and FRQS. We are grateful to F. Bachand, M. Oeffinger and D. Zenkluzen for critical reading of the manuscript and C. Dargemont for helpful discussions. We thank P. Bensidoun and A. Babour for insights about FISH experiments. We also thank A. Fradet-Turcotte, B. Coulombe, D. Finley, M. Kobor, D. Stillman, K. Struhl, F. Winston, P. Hieter and J.M. Huibregtse for sharing reagents.

## AUTHOR CONTRIBUTIONS

Conceptualization, F.R., F.E., and C.J.; Investigation, F.E., C.J., and F.R.; Formal Analysis, F.R.; Writing – Original Draft, F.E.; Writing –Review & Editing, F.R., C.J., F.E., and J.C.; Funding Acquisition, F.R. and J.C.; Supervision, F.R. and J.C.

## ADDITIONAL FILES

**Supplementary file 1**. List of the proteins associated with RNAPII and their differential association in *fcp1-1* cells.

**Supplementary file 2**. List of the proteins associated with Mex67 and their differential association in *ubp15Δ* cells.

**Supplementary file 3**. List of yeast strains used in this study.

**Supplementary file 4**. List of plasmids used in this study.

## DATA AVAILABILITY

Microarray data and processed files have been deposited in GEO under the accession number GSE154671. Mass spectrometry data have been deposited in MassIVE under accession numbers MSV000085729, MSV000085730 and MSV000085731.

---

**Figure 1–figure supplement 1.**
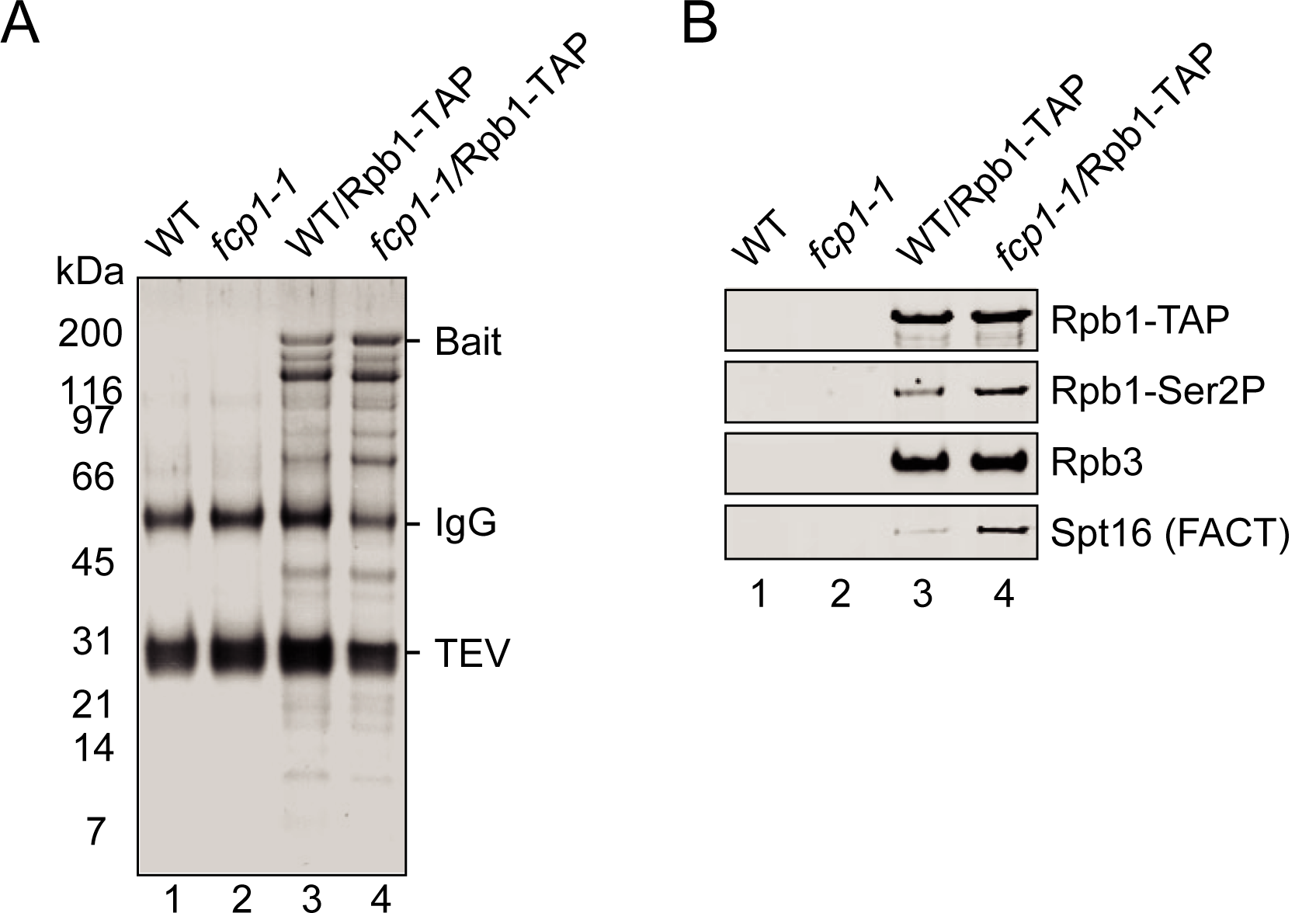
**A**) A silver-stained SDS-PAGE gel showing RNAPII complexes purified from WT and *fcp1-1* cells. Lanes 1 and 2 are control purifications from cells non-expressing Rpb1-TAP protein. The bait (Rpb1-TAP), the IgG (from the beads used to purify the complexes) and the TEV (used to cleave within the TAP-tag and eluate the complexes from the beads) are indicated. **B**) Western blot showing Rpb1, Rpb1-Ser2P, Rpb3 and Spt16 in RNAPII complexes purified from WT and *fcp1-1* cells. Lanes 1 and 2 are control purifications from cells non-expressing Rpb1-TAP protein.

**Figure 2–figure supplement 1.**
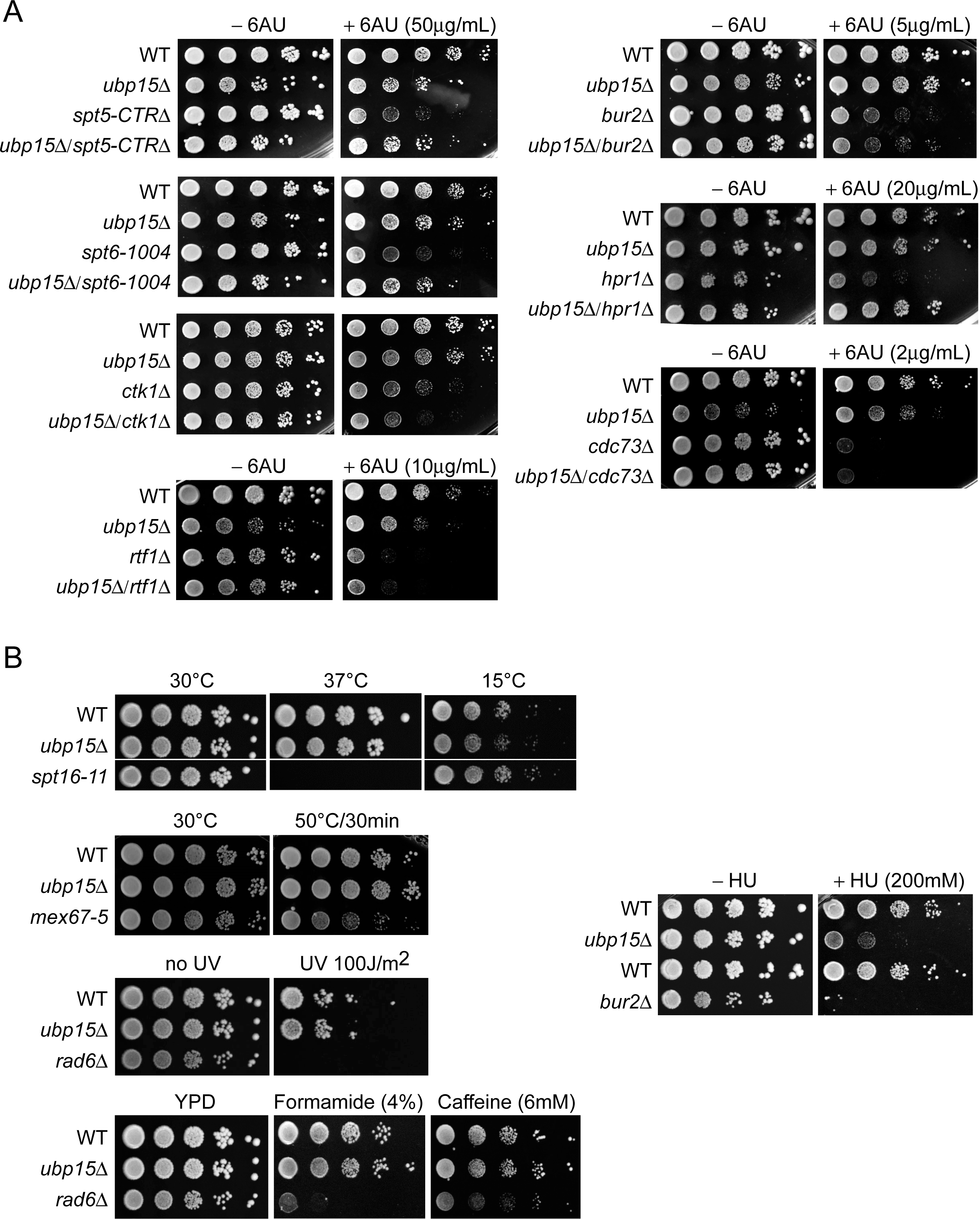
**A**) Deletion of *UBP15* suppresses the 6AU sensitivity of *spt5-CTRΔ, spt6-1004*, and *hpr1Δ* cells but not of *bur2Δ, ctk1Δ, rtf1Δ*, and *cdc73Δ* cells. Cells were grown, diluted and spotted as described for **Figure 2A**. Note that the 6AU concentration varies and has been optimized for each mutant. **B**) Serial-dilution growth assays assessing the sensitivity of WT and *ubp15Δ* cells to various conditions. Cells were grown to saturation in YPD medium, washed, resuspended at the same density in water, serial diluted (fivefold series) and spotted on the indicated media. For growth at 37°C and 15°C, *spt6-11* was used as a positive control. For heat-shock, plates were placed for 30 min at 50°C before incubation at 30°C and *mex67-5* was used as a positive control. For ultraviolet light (UV), the plates were subjected to the indicated doses of UV (UV Stratalinker 1800) right after spotting and *rad6Δ* was used as a positive control. For hydroxyurea (HU), *bur2Δ* was used as a positive control. For formamide and caffeine, *rad6Δ* was used as a positive control.

**Figure 3–figure supplement 1.**
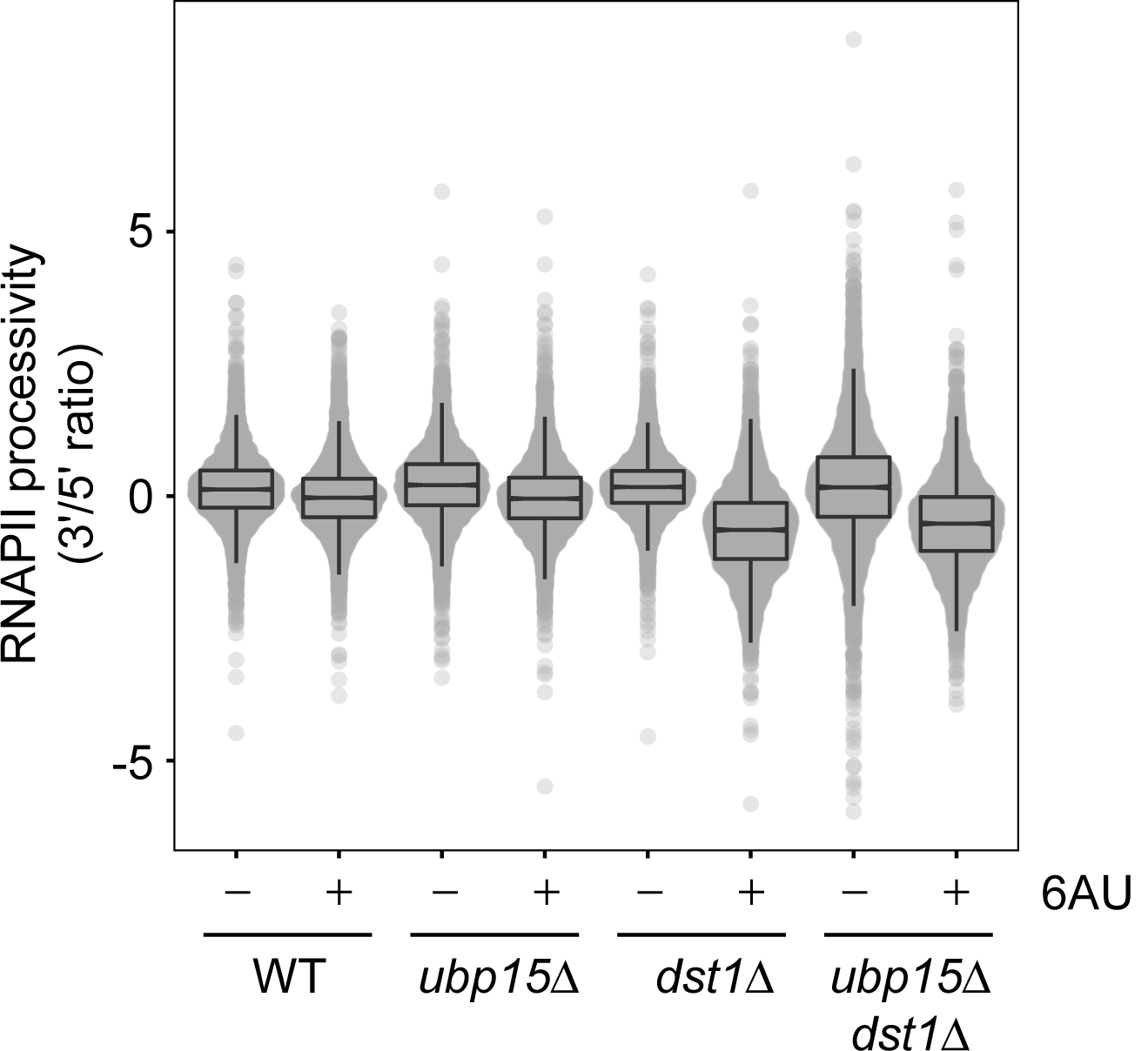
Violin plot showing the RNAPII processivity, as determined by the log_2_ ratio of RNAPII (Rpb3) occupancy in the last 300 bp versus the first 300 bp of each gene, as determined by ChIP-chip. All genes with measurable values are shown (n=4,889).

**Figure 4–figure supplement 1.**
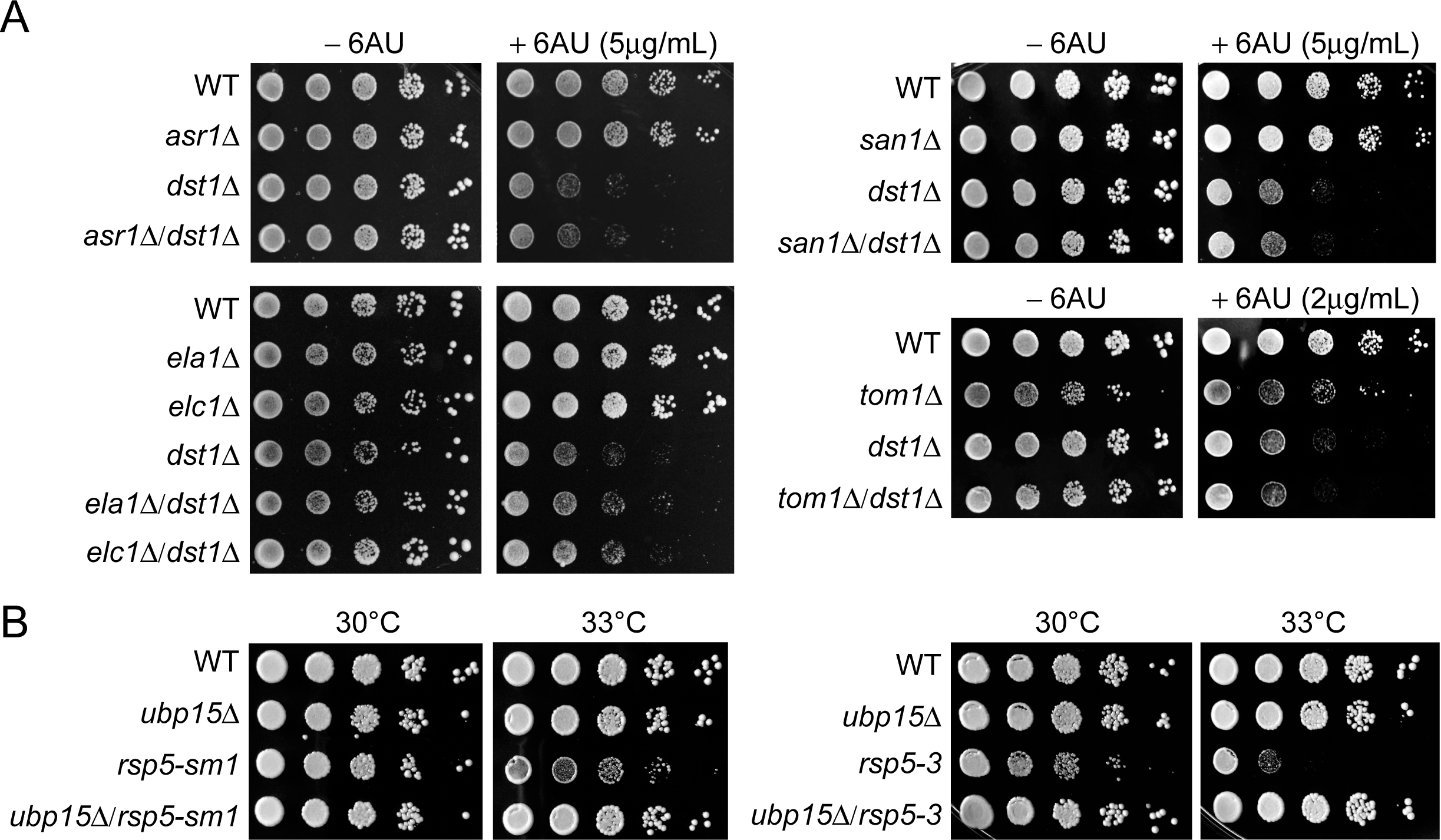
**A**) Serial-dilution growth assays to screen various E3 ligases for a genetic interaction with *DST1*. The indicated yeast strains were grown to saturation in YNB lacking uracil (–URA), washed, resuspended at the same density in water, serially diluted (fivefold series) and spotted on –URA in the absence or presence of 6AU as indicated. Plates were incubated for 3 days at 30°C. **B**) Serial-dilution growth assays assessing the effect of *UBP15* deletion on viability of *rsp5-1, rsp5-sm1* and *rsp5-3* cells at 33°C or 37°C. The indicated yeast strains were grown to saturation in YPD, washed, resuspended at the same density in water, serial diluted (fivefold series) and spotted on YPD. Plates were incubated for 3 days at 30°C or 33°C as indicated.

**Figure 5–figure supplement 1.**
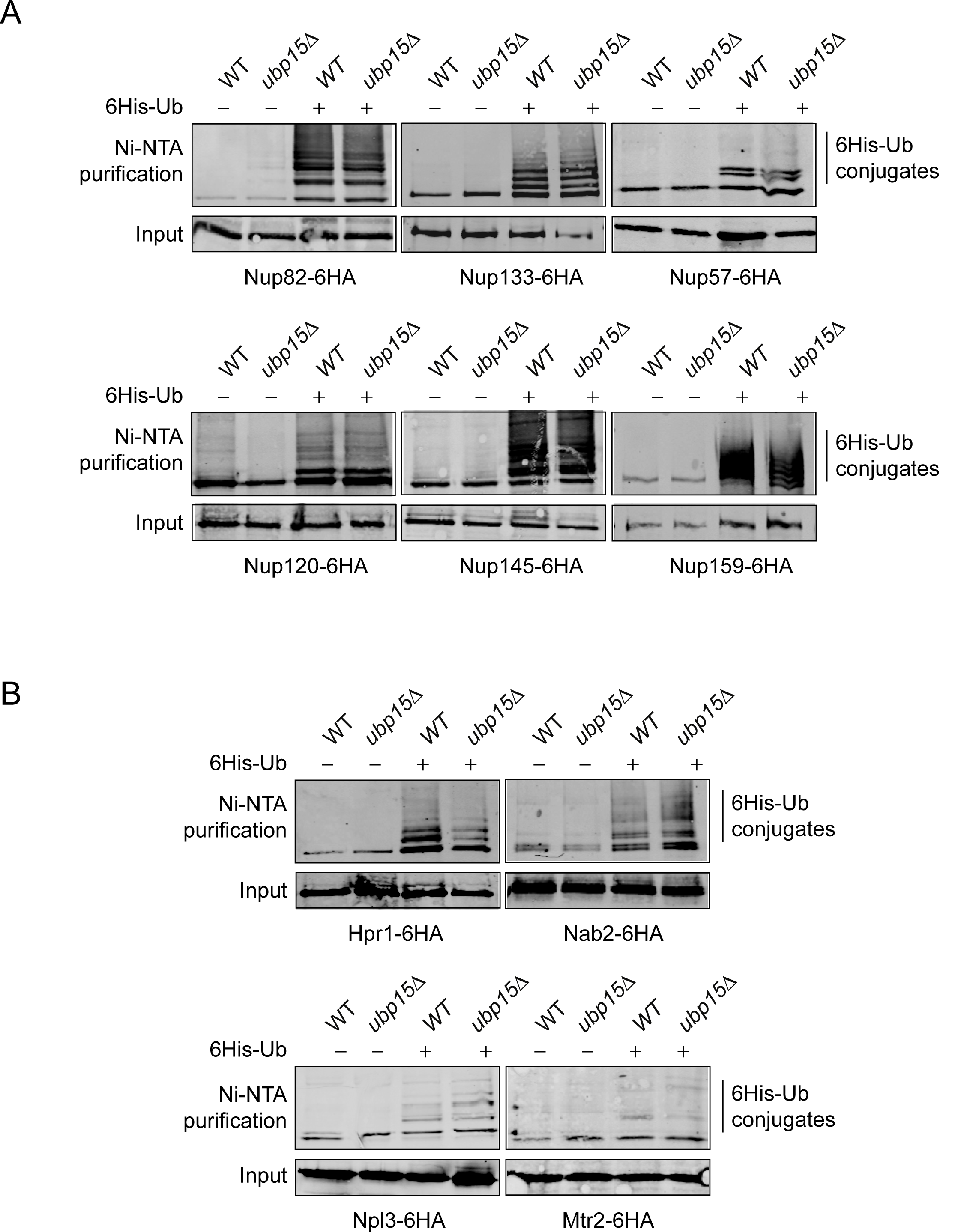
**A**) Western blots for NPC components (Nup82-6HA, Nup133-6HA, Nup57-6HA, Nup120-6HA, Nup145-6HA and Nup159-6HA) levels from His-tagged ubiquitin-conjugated protein pulldowns (Ni-NTA) and their inputs expressing (+) or not (–) 6His-Ub in WT and *ubp15Δ* cells. **B**) Western blots for nuclear export factors (Hpr1-6HA, Nab2-6HA, Npl3-6HA and Mtr2-6HA) levels from His-tagged ubiquitin-conjugated protein pulldowns (Ni-NTA) and their inputs expressing (+) or not (–) 6His-Ub in WT and *ubp15Δ* cells.

**Figure 5–figure supplement 2.**
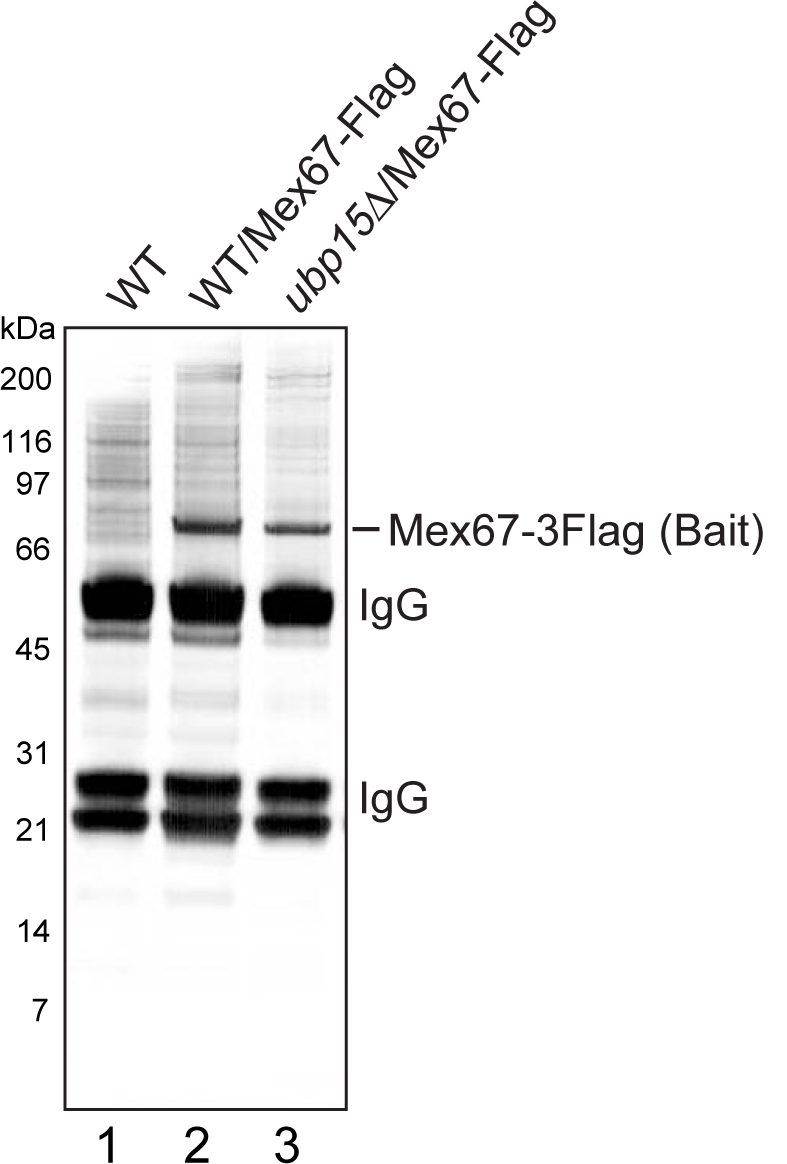
A silver-stained SDS-PAGE gel showing Mex67 complexes purified from WT and *ubp15Δ* cells. Lanes 1 and 2 are control purifications from cells non-expressing any Mex67-3Flag protein. The bait (Mex67-3Flag) and IgG (from the beads used to purify the complexes) are indicated.

## References

Adams RL, Mason AC, Glass L, Aditi, Wente SR. 2017. Nup42 and IP6 coordinate Gle1 stimulation of Dbp5/DDX19B for mRNA export in yeast and human cells. Traffic 18:776–790. https://doi.org/10.1111/tra.12526, PMID: 28869701

Alvarez V, Vinas L, Gallego-Sanchez A, Andres S, Sacristan MP, Bueno A. 2016. Orderly progression through S-phase requires dynamic ubiquitylation and deubiquitylation of PCNA. Sci Rep 6:25513. https://doi.org/10.1038/srep25513, PMID: 27151298

Amerik AY, Li SJ, Hochstrasser M. 2000. Analysis of the deubiquitinating enzymes of the yeast Saccharomyces cerevisiae. Biol Chem 381:981–992. https://doi.org/10.1515/BC.2000.121, PMID: 11076031

Babour A, Dargemont C, Stutz F. 2012. Ubiquitin and assembly of export competent mRNP. Biochim Biophys Acta 1819:521–530. https://doi.org/10.1016/j.bbagrm.2011.12.006, PMID: 22240387

Babour A, Shen Q, Dos-Santos J, Murray S, Gay A, Challal D, Fasken M, Palancade B, Corbett A, Libri D, Mellor J, Dargemont C. 2016. The Chromatin Remodeler ISW1 Is a Quality Control Factor that Surveys Nuclear mRNP Biogenesis. Cell 167:1201–1214 e1215. https://doi.org/10.1016/j.cell.2016.10.048, PMID: 27863241

Bataille AR, Jeronimo C, Jacques PE, Laramee L, Fortin ME, Forest A, Bergeron M, Hanes SD, Robert F. 2012. A universal RNA polymerase II CTD cycle is orchestrated by complex interplays between kinase, phosphatase, and isomerase enzymes along genes. Mol Cell 45:158–170. https://doi.org/10.1016/j.molcel.2011.11.024, PMID: 22284676

Batisse J, Batisse C, Budd A, Bottcher B, Hurt E. 2009. Purification of nuclear poly(A)-binding protein Nab2 reveals association with the yeast transcriptome and a messenger ribonucleoprotein core structure. J Biol Chem 284:34911–34917. https://doi.org/10.1074/jbc.M109.062034, PMID: 19840948

Beaudenon SL, Huacani MR, Wang G, McDonnell DP, Huibregtse JM. 1999. Rsp5 ubiquitin-protein ligase mediates DNA damage-induced degradation of the large subunit of RNA polymerase II in Saccharomyces cerevisiae. Mol Cell Biol 19:6972–6979. https://doi.org/10.1128/mcb.19.10.6972, PMID: 10490634

Beckley JR, Chen JS, Yang Y, Peng J, Gould KL. 2015. A Degenerate Cohort of Yeast Membrane Trafficking DUBs Mediates Cell Polarity and Survival. Mol Cell Proteomics 14:3132–3141. https://doi.org/10.1074/mcp.M115.050039, PMID: 26412298

Brunelle M, Coulombe C, Poitras C, Robert MA, Markovits AN, Robert F, Jacques PE. 2015. Aggregate and Heatmap Representations of Genome-Wide Localization Data Using VAP, a Versatile Aggregate Profiler. Methods Mol Biol 1334:273–298. https://doi.org/10.1007/978-1-4939-2877-4_18, PMID: 26404157

Cabal GG, Genovesio A, Rodriguez-Navarro S, Zimmer C, Gadal O, Lesne A, Buc H, Feuerbach-Fournier F, Olivo-Marin JC, Hurt EC, Nehrbass U. 2006. SAGA interacting factors confine sub-diffusion of transcribed genes to the nuclear envelope. Nature 441:770–773. https://doi.org/10.1038/nature04752, PMID: 16760982

Chapman RD, Heidemann M, Hintermair C, Eick D. 2008. Molecular evolution of the RNA polymerase II CTD. Trends Genet 24:289–296. https://doi.org/10.1016/j.tig.2008.03.010, PMID: 18472177

Cheng H, Dufu K, Lee CS, Hsu JL, Dias A, Reed R. 2006. Human mRNA export machinery recruited to the 5’ end of mRNA. Cell 127:1389–1400. https://doi.org/10.1016/j.cell.2006.10.044, PMID: 17190602

Collin P, Jeronimo C, Poitras C, Robert F. 2019. RNA Polymerase II CTD Tyrosine 1 Is Required for Efficient Termination by the Nrd1-Nab3-Sen1 Pathway. Mol Cell 73:655–669 e657. https://doi.org/10.1016/j.molcel.2018.12.002, PMID: 30639244

Corden JL. 2013. RNA polymerase II C-terminal domain: Tethering transcription to transcript and template. Chem Rev 113:8423–8455. https://doi.org/10.1021/cr400158h, PMID: 24040939

Coulombe C, Poitras C, Nordell-Markovits A, Brunelle M, Lavoie MA, Robert F, Jacques PE. 2014. VAP: a versatile aggregate profiler for efficient genome-wide data representation and discovery. Nucleic Acids Res 42:W485–493. https://doi.org/10.1093/nar/gku302, PMID: 24753414

Daulny A, Geng F, Muratani M, Geisinger JM, Salghetti SE, Tansey WP. 2008. Modulation of RNA polymerase II subunit composition by ubiquitylation. Proc Natl Acad Sci U S A 105:19649–19654. https://doi.org/10.1073/pnas.0809372105, PMID: 19064926

Debelyy MO, Platta HW, Saffian D, Hensel A, Thoms S, Meyer HE, Warscheid B, Girzalsky W, Erdmann R. 2011. Ubp15p, a ubiquitin hydrolase associated with the peroxisomal export machinery. J Biol Chem 286:28223–28234. https://doi.org/10.1074/jbc.M111.238600, PMID: 21665945

Derrer CP, Mancini R, Vallotton P, Huet S, Weis K, Dultz E. 2019. The RNA export factor Mex67 functions as a mobile nucleoporin. J Cell Biol 218:3967–3976. https://doi.org/10.1083/jcb.201909028, PMID: 31753862

Eick D, Geyer M. 2013. The RNA polymerase II carboxy-terminal domain (CTD) code. Chem Rev 113:8456–8490. https://doi.org/10.1021/cr400071f, PMID: 23952966

Fischer T, Rodriguez-Navarro S, Pereira G, Racz A, Schiebel E, Hurt E. 2004. Yeast centrin Cdc31 is linked to the nuclear mRNA export machinery. Nat Cell Biol 6:840–848. https://doi.org/10.1038/ncb1163, PMID: 15311284

Guillemette B, Bataille AR, Gevry N, Adam M, Blanchette M, Robert F, Gaudreau L. 2005. Variant histone H2A.Z is globally localized to the promoters of inactive yeast genes and regulates nucleosome positioning. PLoS Biol 3:e384. https://doi.org/10.1371/journal.pbio.0030384, PMID: 16248679

Gwizdek C, Hobeika M, Kus B, Ossareh-Nazari B, Dargemont C, Rodriguez MS. 2005. The mRNA nuclear export factor Hpr1 is regulated by Rsp5-mediated ubiquitylation. J Biol Chem 280:13401–13405. https://doi.org/10.1074/jbc.C500040200, PMID: 15713680

Gwizdek C, Iglesias N, Rodriguez MS, Ossareh-Nazari B, Hobeika M, Divita G, Stutz F, Dargemont C. 2006. Ubiquitin-associated domain of Mex67 synchronizes recruitment of the mRNA export machinery with transcription. Proc Natl Acad Sci U S A 103:16376–16381. https://doi.org/10.1073/pnas.0607941103, PMID: 17056718

Harlen KM, Churchman LS. 2017. The code and beyond: transcription regulation by the RNA polymerase II carboxy-terminal domain. Nat Rev Mol Cell Biol 18:263–273. https://doi.org/10.1038/nrm.2017.10, PMID: 28248323

Hayakawa A, Babour A, Sengmanivong L, Dargemont C. 2012. Ubiquitylation of the nuclear pore complex controls nuclear migration during mitosis in S. cerevisiae. J Cell Biol 196:19–27. https://doi.org/10.1083/jcb.201108124, PMID: 22213798

Ho HC, MacGurn JA, Emr SD. 2017. Deubiquitinating enzymes Ubp2 and Ubp15 regulate endocytosis by limiting ubiquitination and degradation of ARTs. Mol Biol Cell 28:1271–1283. https://doi.org/10.1091/mbc.E17-01-0008, PMID: 28298493

Hodge CA, Tran EJ, Noble KN, Alcazar-Roman AR, Ben-Yishay R, Scarcelli JJ, Folkmann AW, Shav-Tal Y, Wente SR, Cole CN. 2011. The Dbp5 cycle at the nuclear pore complex during mRNA export I: dbp5 mutants with defects in RNA binding and ATP hydrolysis define key steps for Nup159 and Gle1. Genes Dev 25:1052–1064. https://doi.org/10.1101/gad.2041611, PMID: 21576265

Huibregtse JM, Yang JC, Beaudenon SL. 1997. The large subunit of RNA polymerase II is a substrate of the Rsp5 ubiquitin-protein ligase. Proc Natl Acad Sci U S A 94:3656–3661. https://doi.org/10.1073/pnas.94.8.3656, PMID: 9108033

Hwang GW, Kimura Y, Takahashi T, Lee JY, Naganuma A. 2012. Identification of deubiquitinating enzymes involved in methylmercury toxicity in Saccharomyces cerevisiae. J Toxicol Sci 37:1287–1290. https://doi.org/10.2131/jts.37.1287, PMID: 23208446

Iglesias N, Tutucci E, Gwizdek C, Vinciguerra P, Von Dach E, Corbett AH, Dargemont C, Stutz F. 2010. Ubiquitin-mediated mRNP dynamics and surveillance prior to budding yeast mRNA export. Genes Dev 24:1927–1938. https://doi.org/10.1101/gad.583310, PMID: 20810649

Jani D, Lutz S, Marshall NJ, Fischer T, Kohler A, Ellisdon AM, Hurt E, Stewart M. 2009. Sus1, Cdc31, and the Sac3 CID region form a conserved interaction platform that promotes nuclear pore association and mRNA export. Mol Cell 33:727–737. https://doi.org/10.1016/j.molcel.2009.01.033, PMID: 19328066

Jeronimo C, Bataille AR, Robert F. 2013. The writers, readers, and functions of the RNA polymerase II C-terminal domain code. Chem Rev 113:8491–8522. https://doi.org/10.1021/cr4001397, PMID: 23837720

Jeronimo C, Robert F. 2014. Kin28 regulates the transient association of Mediator with core promoters. Nat Struct Mol Biol 21:449–455. https://doi.org/10.1038/nsmb.2810, PMID: 24704787

Jeronimo C, Watanabe S, Kaplan CD, Peterson CL, Robert F. 2015. The Histone Chaperones FACT and Spt6 Restrict H2A.Z from Intragenic Locations. Mol Cell 58:1113–1123. https://doi.org/10.1016/j.molcel.2015.03.030, PMID: 25959393

Jeronimo C, Collin P, Robert F. 2016. The RNA Polymerase II CTD: The Increasing Complexity of a Low-Complexity Protein Domain. J Mol Biol 428:2607–2622. https://doi.org/10.1016/j.jmb.2016.02.006, PMID: 26876604

Johnson SA, Cubberley G, Bentley DL. 2009. Cotranscriptional recruitment of the mRNA export factor Yra1 by direct interaction with the 3’ end processing factor Pcf11. Mol Cell 33:215–226. https://doi.org/10.1016/j.molcel.2008.12.007, PMID: 19110458

Johnson SA, Kim H, Erickson B, Bentley DL. 2011. The export factor Yra1 modulates mRNA 3’ end processing. Nat Struct Mol Biol 18:1164–1171. https://doi.org/10.1038/nsmb.2126, PMID: 21947206

Kouranti I, McLean JR, Feoktistova A, Liang P, Johnson AE, Roberts-Galbraith RH, Gould KL. 2010. A global census of fission yeast deubiquitinating enzyme localization and interaction networks reveals distinct compartmentalization profiles and overlapping functions in endocytosis and polarity. PLoS Biol 8:e1000471. https://doi.org/10.1371/journal.pbio.1000471, PMID: 20838651

Lee MS, Henry M, Silver PA. 1996. A protein that shuttles between the nucleus and the cytoplasm is an important mediator of RNA export. Genes Dev 10:1233–1246. https://doi.org/10.1101/gad.10.10.1233, PMID: 8675010

Lund MK, Guthrie C. 2005. The DEAD-box protein Dbp5p is required to dissociate Mex67p from exported mRNPs at the nuclear rim. Mol Cell 20:645–651. https://doi.org/10.1016/j.molcel.2005.10.005, PMID: 16307927

Mason PB, Struhl K. 2003. The FACT complex travels with elongating RNA polymerase II and is important for the fidelity of transcriptional initiation in vivo. Mol Cell Biol 23:8323–8333. https://doi.org/10.1128/mcb.23.22.8323-8333.2003, PMID: 14585989

Mason PB, Struhl K. 2005. Distinction and relationship between elongation rate and processivity of RNA polymerase II in vivo. Mol Cell 17:831–840. https://doi.org/10.1016/j.molcel.2005.02.017, PMID: 15780939

McCann TS, Guo Y, McDonald WH, Tansey WP. 2016. Antagonistic roles for the ubiquitin ligase Asr1 and the ubiquitin-specific protease Ubp3 in subtelomeric gene silencing. Proc Natl Acad Sci U S A 113:1309–1314. https://doi.org/10.1073/pnas.1518375113, PMID: 26787877

Mitchell SF, Parker R. 2014. Principles and properties of eukaryotic mRNPs. Mol Cell 54:547–558. https://doi.org/10.1016/j.molcel.2014.04.033, PMID: 24856220

Muratani M, Kung C, Shokat KM, Tansey WP. 2005. The F box protein Dsg1/Mdm30 is a transcriptional coactivator that stimulates Gal4 turnover and cotranscriptional mRNA processing. Cell 120:887–899. https://doi.org/10.1016/j.cell.2004.12.025, PMID: 15797387

Nakazawa Y, Hara Y, Oka Y, Komine O, van den Heuvel D, Guo C, Daigaku Y, Isono M, He Y, Shimada M, Kato K, Jia N, Hashimoto S, Kotani Y, Miyoshi Y, Tanaka M, Sobue A, Mitsutake N, Suganami T, Masuda A, Ohno K, Nakada S, Mashimo T, Yamanaka K, Luijsterburg MS, Ogi T. 2020. Ubiquitination of DNA Damage-Stalled RNAPII Promotes Transcription-Coupled Repair. Cell 180:1228–1244 e1224. https://doi.org/10.1016/j.cell.2020.02.010, PMID: 32142649

Nino CA, Herissant L, Babour A, Dargemont C. 2013. mRNA nuclear export in yeast. Chem Rev 113:8523–8545. https://doi.org/10.1021/cr400002g, PMID: 23731471

Oeffinger M, Wei KE, Rogers R, DeGrasse JA, Chait BT, Aitchison JD, Rout MP. 2007. Comprehensive analysis of diverse ribonucleoprotein complexes. Nat Methods 4:951–956. https://doi.org/10.1038/nmeth1101, PMID: 17922018

Ostapenko D, Burton JL, Solomon MJ. 2015. The Ubp15 deubiquitinase promotes timely entry into S phase in Saccharomyces cerevisiae. Mol Biol Cell 26:2205–2216. https://doi.org/10.1091/mbc.E14-09-1400, PMID: 25877870

Pena A, Gewartowski K, Mroczek S, Cuellar J, Szykowska A, Prokop A, Czarnocki-Cieciura M, Piwowarski J, Tous C, Aguilera A, Carrascosa JL, Valpuesta JM, Dziembowski A. 2012. Architecture and nucleic acids recognition mechanism of the THO complex, an mRNP assembly factor. EMBO J 31:1605–1616. https://doi.org/10.1038/emboj.2012.10, PMID: 22314234

Pepinsky RB. 1991. Selective precipitation of proteins from guanidine hydrochloride-containing solutions with ethanol. Anal Biochem 195:177–181. https://doi.org/10.1016/0003-2697(91)90315-k, PMID: 1888015

Postma M, Goedhart J. 2019. PlotsOfData-A web app for visualizing data together with their summaries. PLoS Biol 17:e3000202. https://doi.org/10.1371/journal.pbio.3000202, PMID: 30917112

Ren B, Robert F, Wyrick JJ, Aparicio O, Jennings EG, Simon I, Zeitlinger J, Schreiber J, Hannett N, Kanin E, Volkert TL, Wilson CJ, Bell SP, Young RA. 2000. Genome-wide location and function of DNA binding proteins. Science 290:2306–2309. https://doi.org/10.1126/science.290.5500.2306, PMID: 11125145

Riles L, Shaw RJ, Johnston M, Reines D. 2004. Large-scale screening of yeast mutants for sensitivity to the IMP dehydrogenase inhibitor 6-azauracil. Yeast 21:241–248. https://doi.org/10.1002/yea.1068, PMID: 14968429

Rodriguez-Navarro S, Fischer T, Luo MJ, Antunez O, Brettschneider S, Lechner J, Perez-Ortin JE, Reed R, Hurt E. 2004. Sus1, a functional component of the SAGA histone acetylase complex and the nuclear pore-associated mRNA export machinery. Cell 116:75–86. https://doi.org/10.1016/s0092-8674(03)01025-0, PMID: 14718168

Rodriguez MS, Gwizdek C, Haguenauer-Tsapis R, Dargemont C. 2003. The HECT ubiquitin ligase Rsp5p is required for proper nuclear export of mRNA in Saccharomyces cerevisiae. Traffic 4:566–575. https://doi.org/10.1034/j.1600-0854.2003.00115.x, PMID: 12839499

Santos-Rosa H, Moreno H, Simos G, Segref A, Fahrenkrog B, Pante N, Hurt E. 1998. Nuclear mRNA export requires complex formation between Mex67p and Mtr2p at the nuclear pores. Mol Cell Biol 18:6826–6838. https://doi.org/10.1128/mcb.18.11.6826, PMID: 9774696

Saroufim MA, Bensidoun P, Raymond P, Rahman S, Krause MR, Oeffinger M, Zenklusen D. 2015. The nuclear basket mediates perinuclear mRNA scanning in budding yeast. J Cell Biol 211:1131–1140. https://doi.org/10.1083/jcb.201503070, PMID: 26694838

Schwabish MA, Struhl K. 2004. Evidence for eviction and rapid deposition of histones upon transcriptional elongation by RNA polymerase II. Mol Cell Biol 24:10111–10117. https://doi.org/10.1128/MCB.24.23.10111-10117.2004, PMID: 15542822

Scott DD, Aguilar LC, Kramar M, Oeffinger M. 2019. It’s Not the Destination, It’s the Journey: Heterogeneity in mRNA Export Mechanisms. Adv Exp Med Biol 1203:33–81. https://doi.org/10.1007/978-3-030-31434-7_2, PMID: 31811630

Sen R, Barman P, Kaja A, Ferdoush J, Lahudkar S, Roy A, Bhaumik SR. 2019. Distinct Functions of the Cap-Binding Complex in Stimulation of Nuclear mRNA Export. Mol Cell Biol 39:e00540–00518. https://doi.org/10.1128/MCB.00540-18, PMID: 30745412

Singh G, Pratt G, Yeo GW, Moore MJ. 2015. The Clothes Make the mRNA: Past and Present Trends in mRNP Fashion. Annu Rev Biochem 84:325–354. https://doi.org/10.1146/annurev-biochem-080111-092106, PMID: 25784054

Stewart M. 2007. Ratcheting mRNA out of the nucleus. Mol Cell 25:327–330. https://doi.org/10.1016/j.molcel.2007.01.016, PMID: 17289581

Strasser K, Bassler J, Hurt E. 2000a. Binding of the Mex67p/Mtr2p heterodimer to FXFG, GLFG, and FG repeat nucleoporins is essential for nuclear mRNA export. J Cell Biol 150:695–706. https://doi.org/10.1083/jcb.150.4.695, PMID: 10952996

Strasser K, Hurt E. 2000b. Yra1p, a conserved nuclear RNA-binding protein, interacts directly with Mex67p and is required for mRNA export. EMBO J 19:410–420. https://doi.org/10.1093/emboj/19.3.410, PMID: 10722314

Strasser K, Masuda S, Mason P, Pfannstiel J, Oppizzi M, Rodriguez-Navarro S, Rondon AG, Aguilera A, Struhl K, Reed R, Hurt E. 2002. TREX is a conserved complex coupling transcription with messenger RNA export. Nature 417:304–308. https://doi.org/10.1038/nature746, PMID: 11979277

Terry LJ, Shows EB, Wente SR. 2007. Crossing the nuclear envelope: hierarchical regulation of nucleocytoplasmic transport. Science 318:1412–1416. https://doi.org/10.1126/science.1142204, PMID: 18048681

Trahan C, Aguilar LC, Oeffinger M. 2016. Single-Step Affinity Purification (ssAP) and Mass Spectrometry of Macromolecular Complexes in the Yeast S. cerevisiae. Methods Mol Biol 1361:265–287. https://doi.org/10.1007/978-1-4939-3079-1_15, PMID: 26483027

Tran EJ, Zhou Y, Corbett AH, Wente SR. 2007. The DEAD-box protein Dbp5 controls mRNA export by triggering specific RNA:protein remodeling events. Mol Cell 28:850–859. https://doi.org/10.1016/j.molcel.2007.09.019, PMID: 18082609

Tufegdzic Vidakovic A, Mitter R, Kelly GP, Neumann M, Harreman M, Rodriguez-Martinez M, Herlihy A, Weems JC, Boeing S, Encheva V, Gaul L, Milligan L, Tollervey D, Conaway RC, Conaway JW, Snijders AP, Stewart A, Svejstrup JQ. 2020. Regulation of the RNAPII Pool Is Integral to the DNA Damage Response. Cell 180:1245–1261 e1221. https://doi.org/10.1016/j.cell.2020.02.009, PMID: 32142654

Tutucci E, Stutz F. 2011. Keeping mRNPs in check during assembly and nuclear export. Nat Rev Mol Cell Biol 12:377–384. https://doi.org/10.1038/nrm3119, PMID: 21602906

Viphakone N, Sudbery I, Griffith L, Heath CG, Sims D, Wilson SA. 2019. Co-transcriptional Loading of RNA Export Factors Shapes the Human Transcriptome. Mol Cell 75:310–323 e318. https://doi.org/10.1016/j.molcel.2019.04.034, PMID: 31104896

Vitaliano-Prunier A, Babour A, Herissant L, Apponi L, Margaritis T, Holstege FC, Corbett AH, Gwizdek C, Dargemont C. 2012. H2B ubiquitylation controls the formation of export-competent mRNP. Mol Cell 45:132–139. https://doi.org/10.1016/j.molcel.2011.12.011, PMID: 22244335

Weirich CS, Erzberger JP, Flick JS, Berger JM, Thorner J, Weis K. 2006. Activation of the DExD/H-box protein Dbp5 by the nuclear-pore protein Gle1 and its coactivator InsP6 is required for mRNA export. Nat Cell Biol 8:668–676. https://doi.org/10.1038/ncb1424, PMID: 16783364

Wende W, Friedhoff P, Strasser K. 2019. Mechanism and Regulation of Co-transcriptional mRNP Assembly and Nuclear mRNA Export. Adv Exp Med Biol 1203:1–31. https://doi.org/10.1007/978-3-030-31434-7_1, PMID: 31811629

Wu X, Chang A, Sudol M, Hanes SD. 2001. Genetic interactions between the ESS1 prolylisomerase and the RSP5 ubiquitin ligase reveal opposing effects on RNA polymerase II function. Curr Genet 40:234–242. https://doi.org/10.1007/s00294-001-0257-8, PMID: 11795843

Xu Z, Wei W, Gagneur J, Perocchi F, Clauder-Munster S, Camblong J, Guffanti E, Stutz F, Huber W, Steinmetz LM. 2009. Bidirectional promoters generate pervasive transcription in yeast. Nature 457:1033–1037. https://doi.org/10.1038/nature07728, PMID: 19169243

Zander G, Hackmann A, Bender L, Becker D, Lingner T, Salinas G, Krebber H. 2016. mRNA quality control is bypassed for immediate export of stress-responsive transcripts. Nature 540:593–596. https://doi.org/10.1038/nature20572, PMID: 27951587

Zenklusen D, Vinciguerra P, Strahm Y, Stutz F. 2001. The yeast hnRNP-Like proteins Yra1p and Yra2p participate in mRNA export through interaction with Mex67p. Mol Cell Biol 21:4219–4232. https://doi.org/10.1128/MCB.21.13.4219-4232.2001, PMID: 11390651

Zenklusen D, Vinciguerra P, Wyss JC, Stutz F. 2002. Stable mRNP formation and export require cotranscriptional recruitment of the mRNA export factors Yra1p and Sub2p by Hpr1p. Mol Cell Biol 22:8241–8253. https://doi.org/10.1128/mcb.22.23.8241-8253.2002, PMID: 12417727

